# PhyloRBT: A Phylogenetic Approach to Detect Reference Bias in Phylogenomic Datasets

**DOI:** 10.64898/2026.07.24.740642

**Authors:** Jeremias Ivan, Robert Lanfear

## Abstract

Recent technological advancements have enabled the rapid generation of high-quality genomes across the tree of life, often resulting in multiple reference genomes for clades of phylogenetic interest. These reference genomes are often used to reconstruct phylogenetically informative loci from short-read data of newly-sequenced species. However, this approach can introduce reference bias where the reconstructed loci have erroneous similarities to those of the reference genome. Since reference bias can seriously affect downstream analyses, it is important to assess its presence in phylogenomic datasets. In this study, we propose PhyloRBT (phylogenetic reference bias test) to detect reference bias by reconstructing each locus multiple times using different reference genomes and then measuring the phylogenetic correlation between these reconstructions and the corresponding locus from the reference genomes. We applied PhyloRBT to hundreds of BUSCO loci reconstructed from short-read data of nine *Eucalyptus* species using 34 different reference genomes. Across the nine species, we found that more than a quarter of the reconstructed loci had significant evidence of reference bias. Compared with the usual approach of mapping to a single closely-related reference genome and using all reconstructed loci, excluding the putatively biased loci also produced a species tree topology that is more consistent with the published tree. In conclusion, PhyloRBT offers a straightforward way to detect reference bias in individual loci, and to selectively remove those biased loci from downstream analyses.

## Introduction

Modern phylogenomic studies use hundreds or thousands of loci to infer various aspects of the evolutionary history of many taxonomic groups (Young and Gillung 2020; Lozano-Fernandez 2022). For example, He et al. (2024) used 1,764 single-copy, complete BUSCO loci (Simão et al. 2015) from 487 Basidiomycota species to infer the species tree and estimate the divergence times of the group, while Stiller et al. (2024) estimated the phylogeny of 363 Neoaves (modern birds) species using 63,430 intergenic loci. While these inferences aimed to answer different evolutionary questions using different approaches, all rely on the accuracy of individual gene sequences and gene trees being analysed. When a high-quality reference genome is not available for the species of interest, locus sequences are often reconstructed by mapping the sequencing reads of the newly-sequenced species to one or more reference genomes from closely-related taxa (e.g., Croucher et al. 2011; Bayly et al. 2013; Pease et al. 2016; Edelman et al. 2019). However, mapping can introduce *reference bias,* where the sequencing reads from the newly-sequenced species are more likely to map successfully to a reference genome when they contain variants that make them more similar to that genome, while reads with non-reference alleles might be mapped to the wrong region of the genome or not mapped at all (Degner et al. 2009; Stevenson et al. 2013; Lin et al. 2024). As a result, reconstructed sequences that are derived from the mapped reads can be erroneously similar to the reference genome. Reference bias has been shown to affect population genetic inferences (Nevado et al. 2014; Brandt et al. 2015; Gopalakrishnan et al. 2017; Shafer et al. 2017; Günther and Nettelblad 2019), quantification of gene expression (Munger et al. 2014; Panousis et al. 2014; Zhan et al. 2021; Coombes et al. 2024), and phylogenetic inferences. For example, Valiente-Mullor et al. (2021) showed that the positions of two *Legionella pneumophila* isolates tended to cluster together with their reference genomes on the species tree, while Bertels et al. (2014) showed that using a single reference genome during mapping can result in an inaccurate tree especially when analysing highly divergent taxa. This also highlights that reference bias can be exacerbated when the reference and the newly-sequenced species are more evolutionarily diverged (Gopalakrishnan et al. 2017; Bohling 2020; Huang et al. 2024). As reference bias can seriously impact downstream analyses, it is important to assess its presence in empirical datasets.

Various approaches have been developed to assess reference bias in genomic datasets. For example, AMBER measures reference bias on ancient DNA datasets based on metrics such as mapping breadth and depth (Dolenz et al. 2024). Similarly, `biastools` computes a bias score for diploid individuals based on read depth, density of the alternative alleles, and frequency of sites that are inconsistent with the diploid state (Lin et al. 2024). Another approach that has been suggested to mitigate reference bias is to use multiple reference genomes (Bertels et al. 2014; Chen et al. 2021) or pan-genome graphs (Schneeberger et al. 2009; Valenzuela et al. 2018; Kim et al. 2019; Sirén et al. 2021) from closely-related species during mapping. This is becoming increasingly common in phylogenomic studies because long read sequencing has enabled *de novo* assembly of high-quality reference genomes for many groups, including non-model organisms (Ekblom and Wolf 2014; Sedlazeck et al. 2018; Rhie et al. 2021). In principle, the availability of multiple reference genomes should minimise the effects of reference bias, because they should collectively better represent the allelic variation in the newly-sequenced species. It also presents an opportunity to measure reference bias in new ways.

In this study, we propose PhyloRBT (phylogenetic reference bias test), a new approach that leverages the availability of multiple reference genomes to measure reference bias in individually reconstructed loci from short-read data. PhyloRBT works by first mapping the short reads from a newly-sequenced species to multiple reference genomes. Then, we reconstruct each locus of interest once for each reference genome. In an ideal world (i.e., when there is no reference bias or other errors in the reconstruction), the sequence of each individual locus should be identical no matter which reference genome was used during its reconstruction. However, in the presence of reference bias, the sequence of an individual locus will tend to be more similar to the same locus in the reference genome used to reconstruct it than would be expected by chance. We leverage this difference using statistical tests to detect reference bias (see MATERIALS AND METHODS section for details). In this study, we focus on BUSCO (Benchmarking Universal Single Copy Orthologs; Simão et al. 2015; Manni et al. 2021) as our target loci because they are widely-used for phylogenomic inference (e.g., Kanzi et al. 2020; Li et al. 2021; Ferguson et al. 2024a) and expected to be present in single copies in most genomes, which minimises the effects of paralogs in our analyses.

To infer BUSCO loci from the short-read data of a newly-sequenced species, researchers need to first assemble the reads before running the BUSCO pipeline (Simão et al. 2015), which can be done in two ways. One option is to do *de novo* assembly of the short-read data. This avoids reference bias altogether because no reference genomes are used during the assembly. However, *de novo* assembly from exclusively short reads can be challenging, as these assemblies tend to be more fragmented and contain a higher proportion of gaps compared to those built from long sequencing reads (Paszkiewicz and Studholme 2010; Liao et al. 2019). An alternative and commonly-used approach involves mapping the short reads to a closely-related reference genome, followed by variant calling to generate a consensus sequence. This method is susceptible to reference bias as the reconstructed BUSCO sequences may be affected by how the short sequencing reads align to the reference genome in the first place. Here, we focus on the second approach (i.e., mapping) to infer BUSCO loci from short-read datasets and assess how reference bias may affect their sequence reconstruction.

In this study, we first describe PhyloRBT to detect reference bias in phylogenomic datasets. We then apply PhyloRBT to assess the extent of reference bias in an empirical dataset from eucalypts, comprising *Eucalyptus*, *Corymbia*, and *Angophora*. Specifically, we analyse the dataset from Ferguson et al. (2024a) because: (i) it consists of high-quality reference genomes and their associated short sequencing reads from 35 species, which enables us to compare the reconstructed loci (inferred from the mapped reads) with locus sequences inferred independently from reference genomes assembled *de novo*, such that the latter cannot suffer from reference bias; (ii) the 35 reference genomes come from different parts of the eucalypt species tree (Crisp et al. 2024; Ivan et al. 2026), allowing us to test different scenarios where the reference genomes come from closely or distantly-related taxa; (iii) eucalypt genomes have been shown to exhibit high heterozygosity and structural rearrangements (Grattapaglia et al. 2011; Silva-Junior and Grattapaglia 2015; Ferguson et al. 2024a, 2024b), which means that this dataset might be more sensitive to the selection of reference genomes during mapping and is therefore useful to clearly demonstrate the effects of reference bias in phylogenomic inference; and (iv) an independently-inferred species tree was available for this group (Ferguson et al. 2024a), allowing us to assess the downstream impact of reference bias on species tree inference. To test our method, we selected the short-read data from nine *Eucalyptus*species that are spread across the species tree of the group (Ferguson et al. 2024a) and individually mapped each to all 34 taxa in the dataset that are not from the same species as the short-read data. We show that statistically significant evidence of reference bias is found in a large proportion of the reconstructed BUSCO loci across species, and describe how this can affect the downstream phylogenomic inferences.

## Materials and Methods

### Overview of PhyloRBT

In order to measure the extent of reference bias in phylogenomic datasets, we designed PhyloRBT that leverages the availability of multiple reference genomes and focuses on one locus at a time. All the code necessary to reproduce the methods in this paper is available at https://github.com/jeremiasivan/PhyloRBT (DOI: 10.5281/zenodo.23328980).

PhyloRBT works by using each reference genome in turn to reconstruct each locus for each set of sequencing reads. For example, if there are three reference genomes available for the clade of interest, we would reconstruct each locus three times (once for each reference genome) for each newly-sequenced species in our dataset (where each new species is represented by a single set of short sequencing reads). If there is no reference bias and no error in the reconstructions, the three reconstructed loci from the same set of sequencing reads would be identical to each other. In contrast, reference bias would make the three reconstructed sequences more similar to the reference genomes than we would expect by chance. This pattern is different from the stochastic error that may occur during the mapping and variant calling processes, which would make the three reconstructed locus sequences different from one another independently of the reference genome used to reconstruct them. PhyloRBT distinguishes reference bias from stochastic error by building a phylogenetic tree of all reconstructed versions of the locus of interest along with the sequences of the same locus extracted from the reference genomes. We then ask if the *reconstructed* loci are more similar to their corresponding *reference* loci than would be expected by chance, indicating evidence of reference bias.

Figure 1 gives an example where we are interested in reconstructing a single locus from a single set of short reads (S1), and we have three reference genomes available from the clade of interest (*RefA*, *RefB*, *RefC*). First, we reconstruct three versions of the respective locus for our target species (S1) by mapping the short reads to each of the three reference genomes. We then build a locus tree from the three versions of the reconstructed locus, as well as the locus sequences from the three reference genomes, resulting in a tree with 6 tips. In a situation without reference bias, the three reconstructed loci from S1 would be identical to one another, and the locus tree would group the three reconstructed loci together (i.e., *S1--RefA, S1--RefB, and S1--RefC* are grouped together on Fig. 1a). In the presence of reference bias, the reconstructed locus from S1 would cluster with the locus sequence from the reference genome used to reconstruct it (i.e., *S1--RefA* groups with *RefA*, *S1--RefB* groups with *RefB*, and *S1--RefC* groups with *RefC* on Fig. 1b). The stronger the effect of reference bias, the stronger the clustering of the reconstructed loci with their respective reference loci.

**Figure 1.**
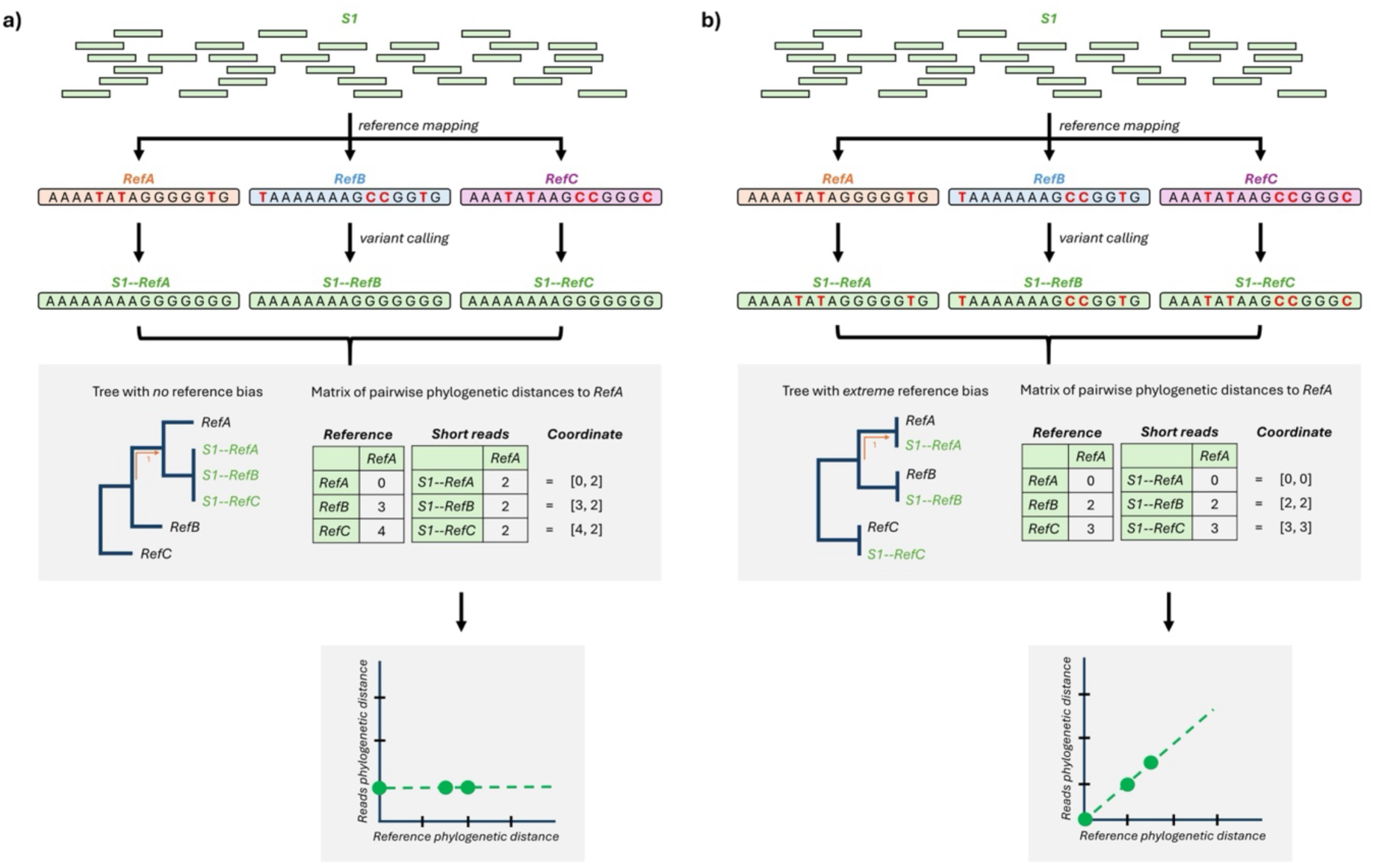
Overview of PhyloRBT when there is (a) no reference bias, and (b) extreme reference bias. Pairwise phylogenetic distance refers to the pairwise distance in branch length units. S1 denotes a single set of short reads from a newly-sequenced species, while the *S1--RefA/B/C* notation refers to the short reads from S1 that are mapped to the reference genome from species A/B/C. Colouring is based on the species.

PhyloRBT tests for the presence of reference bias on a given locus by comparing the phylogenetic distances of the reconstructed loci to their respective reference loci. In order to do this, we start by arbitrarily choosing a single sequence from the reference genomes as the focal sequence to avoid pseudoreplication (e.g., *RefA* on Fig. 1). Then, we measure the pairwise phylogenetic distance of each reference locus to the focal tip (i.e., reference phylogenetic distance, x-axis on Fig. 1), as well as the pairwise phylogenetic distance of each reconstructed locus to the focal tip (i.e., reads phylogenetic distance, y-axis on Fig. 1). We use a linear model and a Spearman’s rank order correlation test to ask if the two measures of pairwise phylogenetic distances are more correlated than would be expected by chance (Schober et al. 2018). The linear model measures the strength of linear association (where *x* = *y* reflects an extreme reference bias; Fig. 1), while Spearman’s rank order correlation test does not assume normality and is robust to non-linear association and outliers. As a result, using both tests would provide a more robust detection of reference bias in individual loci. Significant results (*p*<0.05) from either or both tests would suggest the presence of reference bias for a single locus reconstructed from a single set of short reads that are mapped to multiple references.

### Application to an Empirical Dataset

#### Overview

We applied PhyloRBT to an empirical dataset of 35 eucalypts (comprising *Eucalyptus*, *Corymbia*, and *Angophora*), where each species has its own well-polished *de novo* reference genome and corresponding set of short reads (Ferguson et al. 2024a). We used short-read data from nine representative *Eucalyptus* species, which were individually mapped to 34 reference genomes. Then, we ran the BUSCO pipeline to extract BUSCO loci from each set of mapped reads, resulting in 34 reconstructions of each BUSCO locus for each of the nine short-read species in our dataset. We then used PhyloRBT to test for reference bias in each BUSCO locus from each species, and assessed whether removing BUSCO loci with evidence of reference bias affected the species tree inference of the group.

#### Datasets

Our dataset comprises 35 taxa from eucalypts: 32 *Eucalyptus*, 2 *Corymbia* (*C. maculata*, *C. calophylla*), and one *Angophora* (*A. floribunda*) (Ferguson et al. 2024a). We selected nine representative *Eucalyptus* taxa to test our method – this should be sufficient to effectively test our new approach and evaluate the scale of reference bias in a dataset, but small enough to ensure computational efficiency. The nine representative taxa represent three groups of three closely-related species that are spread across the species tree from Ferguson et al. (2024a): *E. curtisii*, *E. erythrocorys*, and *E. tenuipes* in group one; *E. grandis*, *E. globulus*, and *E. viminalis* in group two; and *E. albens*, *E. melliodora*, and *E. sideroxylon* in group three (Fig. S1). This configuration allowed us to test the presence of reference bias in BUSCO inference when a set of short reads was mapped to a reference coming from closely-related taxa (e.g., *E. curtisii* short reads mapped to *E. tenuipes* reference genome), distantly-related taxa (e.g., *E. curtisii* short reads mapped to *E. albens* reference genome), and different genera (e.g., *E. curtisii* short reads mapped to *A. floribunda* reference genome). To avoid circularity, short reads were never mapped to the genome from the same species from which they were derived (see below).

#### Reference Genomes: Downloading Data and Extracting Reference BUSCO

We downloaded the reference genomes of the 35 eucalypt species from NCBI (BioProject PRJNA509734) using NCBI Datasets (https://www.ncbi.nlm.nih.gov/datasets/) based on their accession numbers. In order to extract BUSCO from the reference genomes, we ran BUSCO pipeline v5.8.0 (Manni et al. 2021) on each reference genome using `eudicots_odb10` as the lineage dataset and MetaEuk (Levy Karin et al. 2020) as the gene predictor tool. We chose *eudicots* because it is the most specific taxonomic group that encompasses eucalypts, while MetaEuk provides a balance between speed (i.e., it is faster than AUGUSTUS (Stanke et al. 2004)) and accuracy (i.e., it is slower than miniprot (Li 2023) but less sensitive when the assemblies are highly-divergent from the database). We then retrieved all BUSCO that were identified as single-copy and complete for each of the 35 reference genomes, excluding duplicated and fragmented BUSCO to ensure that the subsequent phylogenetic inferences were not affected by paralogs. The code to run this analysis is available at https://github.com/jeremiasivan/PhyloRBT/tree/main/data_preparation/2_extract_busco.

#### Short Reads

### Downloading raw data, quality control, and mapping

We downloaded the raw short sequencing reads of the nine representative *Eucalyptus* taxa mentioned above from NCBI (BioProject PRJNA509734) using SRA Toolkit (https://github.com/ncbi/sra-tools) based on their accession numbers, consisting of two files per taxon: forward and reverse reads. Then, we performed quality control on each set of short reads using AdapterRemoval v2 (Schubert et al. 2016) by trimming the adapters and filtering out low-quality reads (Q<25) as well as leading and trailing ‘N’ using the command:

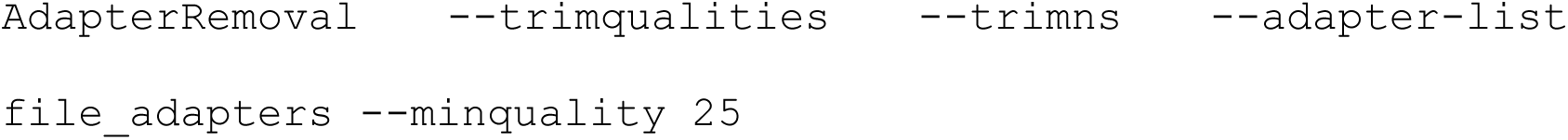

In order to mimic an empirical scenario in which a reference genome from the same species as the newly-sequenced species is unavailable, we mapped the short-read data of each of the nine representative *Eucalyptus* species to all reference genomes except the one from the same species. To do this, we used BWA-MEM2 (Li and Durbin 2009; Vasimuddin et al. 2019) with default settings, resulting in 34 SAM files per representative species. We compressed the SAM files to BAM format and sorted the mapped reads based on their genomic coordinates using SAMtools v1.18 (Li et al. 2009). Then, we removed putative PCR duplicates using SAMtools `rmdup` (Li et al. 2009) and performed variant calling using `mpileup` command from BCFtools v1.19 (Li 2011). Lastly, we filtered out low-quality variants (`QUAL`≤`15` or `MAPQ`≤`30`) (Ferguson et al. 2024b), did variant normalisation, and generated the consensus sequence for each BAM file using BCFtools `view`, `norm`, and `consensus`, respectively, totalling 306 consensus sequences. The code to run this analysis is available at https://github.com/jeremiasivan/PhyloRBT/tree/main/data_preparation/1_readmap.

### Extracting BUSCO

In order to extract BUSCO from the consensus sequences, we ran the BUSCO pipeline on each consensus sequence with the same parameters as the ones used for the reference genomes. Then, for each of the nine representative species, we retrieved all BUSCO that were found as single-copy and complete in *all* versions of the consensus sequences – representing different reference genomes used during mapping – and *all* 34 reference genomes being used. Moreover, in order to ensure that each reconstructed BUSCO from the consensus sequence is well-supported by the short reads (i.e., if the BUSCO has zero coverage on the consensus sequence, then the reconstructed BUSCO would only reflect the sequence of the reference genome), we measured the coverage of each BUSCO on each consensus sequence by first converting the BUSCO GFF files outputted by the BUSCO pipeline to a BED format using BEDOPS v2.4.41 (Neph et al. 2012). Then, we calculated the coverage of each BUSCO on each consensus sequence using SAMtools `depth` (Li et al. 2009). We set the minimum coverage for each BUSCO to be 10x for *all* consensus sequences (because the mean sequencing depth of the short reads is ∼20x), while the maximum is two times the mean coverage of the mapped reads (Table S1). These are strict filters that would not be necessary for a typical phylogenomic analysis. However, they are helpful to accurately assess the scale of reference bias in the dataset because every reconstructed locus is present and well-supported for every species, ensuring that missing data does not affect our power to detect reference bias in this study. The code to run this analysis is available at https://github.com/jeremiasivan/PhyloRBT/blob/main/data_preparation/2_extract_busco/3_shortreads.Rmd.

#### Generating BUSCO Alignments and Trees

For each BUSCO from each representative species, we extracted its amino acid sequences across all 34 consensus sequences, which represent different reference genomes. We then combined these sequences, as well as BUSCO amino acid sequences from all reference genomes, into one FASTA file (i.e., each FASTA file contained 34 reconstructed versions of the locus from the representative species, as well as the 34 versions of the same locus that were extracted directly from the reference genomes). We then ran MAFFT v7.520 (Katoh and Standley 2013) with default settings (FFT-NS-2) to align these 68 sequences. In order to assess the effect of filtering out poorly-aligned regions from the alignments on reference bias, we generated another set of BUSCO alignments where each alignment was filtered using the `-automated1` flag from trimAl v1.2rev59 (Capella-Gutiérrez et al. 2009). We then built phylogenies for the trimmed and untrimmed alignments using IQ-TREE2 (Minh et al. 2020b) with 1,000 UFBoot replicates (Hoang et al. 2018) and the best model from ModelFinder (Kalyaanamoorthy et al. 2017), resulting in two gene trees (based on the filtered and unfiltered alignments) for each BUSCO locus. The code to run this analysis is available at https://github.com/jeremiasivan/PhyloRBT/blob/main/data_preparation/2_extract_busco/4_gene_trees.Rmd.

#### Measuring Reference Bias

We ran PhyloRBT on individual BUSCO trees to detect reference bias on each tree (from both the trimmed and untrimmed alignments). PhyloRBT first calculates the pairwise phylogenetic distances between all the reference genomes and consensus sequences using `cophenetic.phylo` function in ape (Paradis et al. 2004). In order to measure reference bias for each set of short reads in each tree, we arbitrarily selected the locus from the reference genome of *E. cloeziana* as our focal sequence (note that the selection of focal species should not affect the results; Fig. 1), then extracted the pairwise distances from the focal sequence to each reference genome and to each reconstructed sequence of the representative species (see Fig. 1 for reference). Finally, we ran a linear model and a Spearman’s rank order correlation test to test for significant associations between the two sets of distances using R (R Core Team 2023), and counted the proportion of BUSCO loci with statistically significant results (*p*<0.05) for each test independently, and the proportion when both tests were significant. The code to run this analysis is available at https://github.com/jeremiasivan/PhyloRBT/blob/main/codes/2_run.Rmd.

#### Validating PhyloRBT against Reference-Bias-Free Sequences

For each BUSCO from each representative species, we retrieved its amino acid sequences from the *Generating BUSCO Alignments and Trees* step above. In addition, we added the BUSCO sequence inferred from the *de novo* (i.e., reference-bias free) reference genome that comes from the same species as the short-read data, resulting in 69 sequences per BUSCO (34 reconstructions + 35 reference genomes). If a BUSCO was not found to be single-copy or complete in the *de novo* reference genome, it was excluded from the subsequent analyses. We then aligned each BUSCO with MAFFT (Katoh and Standley 2013) with the `--auto` flag. For each BUSCO alignment, we then compared each reconstruction to: (i) the reference it was mapped to, and (ii) the *de novo* reference and counted the following: `n_sites` (i.e., total number of sites, including gaps and unknown amino acids), `n_valid` (i.e., number of sites, excluding gaps and unknown amino acids in any sequence), `n_divergent` (i.e., number of sites that differ between the two reference genomes), `n_biased` (i.e., number of `n_divergent` sites where the reconstruction follows the reference it was mapped to), and `n_other` (i.e., number of `n_valid` sites where the reconstruction does not follow either reference).

To assess the predictive power of PhyloRBT, we calculated the sum proportion of reference bias across reconstructions per locus (Σ*_n_*_.*biased*_ / Σ*_n_*_.*valid*_), sorted the loci from the one with the highest proportion of bias, and calculated the percentage of the top-N most-biased loci that were among the N loci flagged by PhyloRBT. We also repeated the linear model and Spearman’s rank order correlation test for every BUSCO using the amino-acid p-distances: the reads distance was the p-distance between each reconstruction and the *de novo* reference sequence ((`n_biased` + `n_other`) / `n_valid`), while the reference distance was the p-distance between the reference genome it was mapped to and the *de novo* reference sequence (`n_divergent` / `n_valid`) (see Figure 1 for reference). We then calculated the proportion of loci with significant results based on different tests and compared it with the PhyloRBT results.

#### Assessing the Effect of Reference Bias using Concordance Factors

In the absence of reference bias, *all* reconstructions of the same sample (or species) should cluster together on the tree (Fig. 1). In order to assess the utility of PhyloRBT to detect putatively biased loci and improve the clustering of the species reconstructions by excluding such loci, we inferred ASTRAL trees using ASTRAL-III (Zhang et al. 2018) for each of the nine representative species (with the 34 reference genomes) based on three sets of BUSCO gene trees: (i) all BUSCO loci, (ii) BUSCO with significant evidence of reference bias from *either* or both statistical tests, and (iii) BUSCO with *no* evidence of reference bias based on the statistical tests. Here, we filtered out loci with significant results from *either* test because this criterion represents the most conservative approach to minimise reference bias in the dataset (but also the most aggressive approach that removes the most loci). We hypothesise that the clustering of the newly-sequenced species would have higher concordance factors (CFs) on the ASTRAL tree that was constructed only from putatively unbiased loci compared to when using all loci.

For each of the 27 ASTRAL trees (i.e., 9 representative species × 3 sets of BUSCO trees), we retrieved the ASTRAL posterior probability of the branch that unites *all* 34 reconstructions of the newly-sequenced species. We also retrieved the ASTRAL quartet score (qCF) and computed the gCF (gene concordance factor) and sCF (site concordance factor) of the same branch as these metrics are designed to measure the variation of tree topologies within the data (i.e., how many quartets, genes, and individual sites support that branch) (Minh et al. 2020a; Mo et al. 2023; Lanfear and Hahn 2024). We computed the gCF and sCF of each ASTRAL tree using IQ-TREE2 (Minh et al. 2020b) with the following commands:

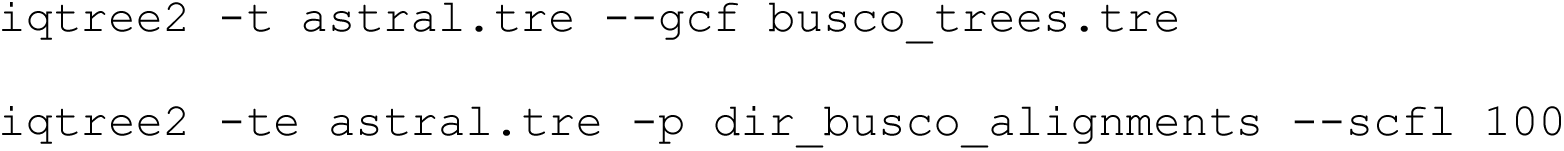

where `astral.tre` reflects the ASTRAL tree, `busco_trees.tre` refers to a file containing the list of BUSCO trees in Newick format with one tree per line, and `dir_busco_alignments` refers to the directory where individual BUSCO alignments are stored. The code to run this analysis is available at https://github.com/jeremiasivan/PhyloRBT/blob/main/codes/3_topology_test.Rmd.

#### Assessing the Effect of Reference Bias on Species Tree Inference

While including all reconstructions of the same species on the phylogeny is informative to measure the support of the clustering, most phylogenomic studies would only use one reconstruction of the newly-sequenced species on the species tree. In order to assess the effect of reference bias on individual reconstructions, we first selected one species from each group (*E. albens*, *E. erythrocorys*, and *E. grandis*; Fig. S1). For each representative species, we then built ASTRAL trees that consist of one reconstruction of the species and the 34 reference genomes using two sets of BUSCO loci: *all* BUSCO and only putatively unbiased BUSCO, totalling 204 ASTRAL trees (3 species × 34 reconstructions × 2 sets of BUSCO trees). However, we constrained the tree topology of the 34 reference genomes using the published tree (Ferguson et al. 2024a) in ASTRAL v5.6.9 (Rabiee and Mirarab 2020), which allows us to compare the placement of the newly-sequenced species on the tree when mapped to different reference genomes.

For each reconstruction, we then assessed if excluding the putatively biased loci from the ASTRAL tree inference improved the consistency of the tree compared to the published tree (Ferguson et al. 2024a). We measured this by calculating the phylogenetic distance (in coalescent units) of the species placement between the two trees – namely how many coalescent units does the species need to traverse moving from its placement on the ASTRAL tree to reach its placement (i.e., branch) on the published tree. We hypothesise that the ASTRAL tree inferred from only putatively unbiased BUSCO would have lower phylogenetic distance (to the published tree) than the one inferred from all BUSCO. The code to run this analysis is available at https://github.com/jeremiasivan/PhyloRBT/blob/main/codes/3_topology_test.Rmd.

## Results

### Reference bias is pervasive in phylogenomic datasets

Across the nine representative species, the average coverage of mapped short reads ranged from 8.214x (for *E. tenuipes* mapped to *C. maculata*) to 37.564x (for *E. sideroxylon* mapped to *E. tenuipes*), with generally lower coverage and percentage of mapped reads when using *Corymbia* and *Angophora* reference genomes (Tables S1-S2). For each of the nine short-read datasets, our analyses resulted in the reconstruction of between 192 and 466 BUSCO loci (Table S3). Based on *both* the linear model and Spearman’s rank order correlation test, approximately 42-49% (average 45.53%) of these loci show significant evidence of reference bias across the representative species (Fig. 2a, Table S3). This proportion is even higher when considering individual tests: 66-75% (average 70.11%) based on the linear model only, 48-57% (average 51.68%) based on the Spearman’s rank order correlation test only, and 73-83% (average 76.27%) based on *either* test (Fig. 2a, Table S3). When excluding the distantly-related *Angophora* and *Corymbia* reference genomes from the analyses, the proportion of BUSCO loci with significant results decreases to 25-31% (average 28.23%) for *both* tests combined; 46-54% (average 50.66%) for only linear model; 32-37% (average 34.58%) for only Spearman’s rank order correlation test, and 52-62% (average 57.01%) for *either* test (Fig. 2b, Table S3). Filtering out poorly-aligned regions made negligible and inconsistent changes to the proportion of BUSCO loci with significant evidence of reference bias across representative species (Table S4), so for brevity we do not discuss it further here.

**Figure 2.**
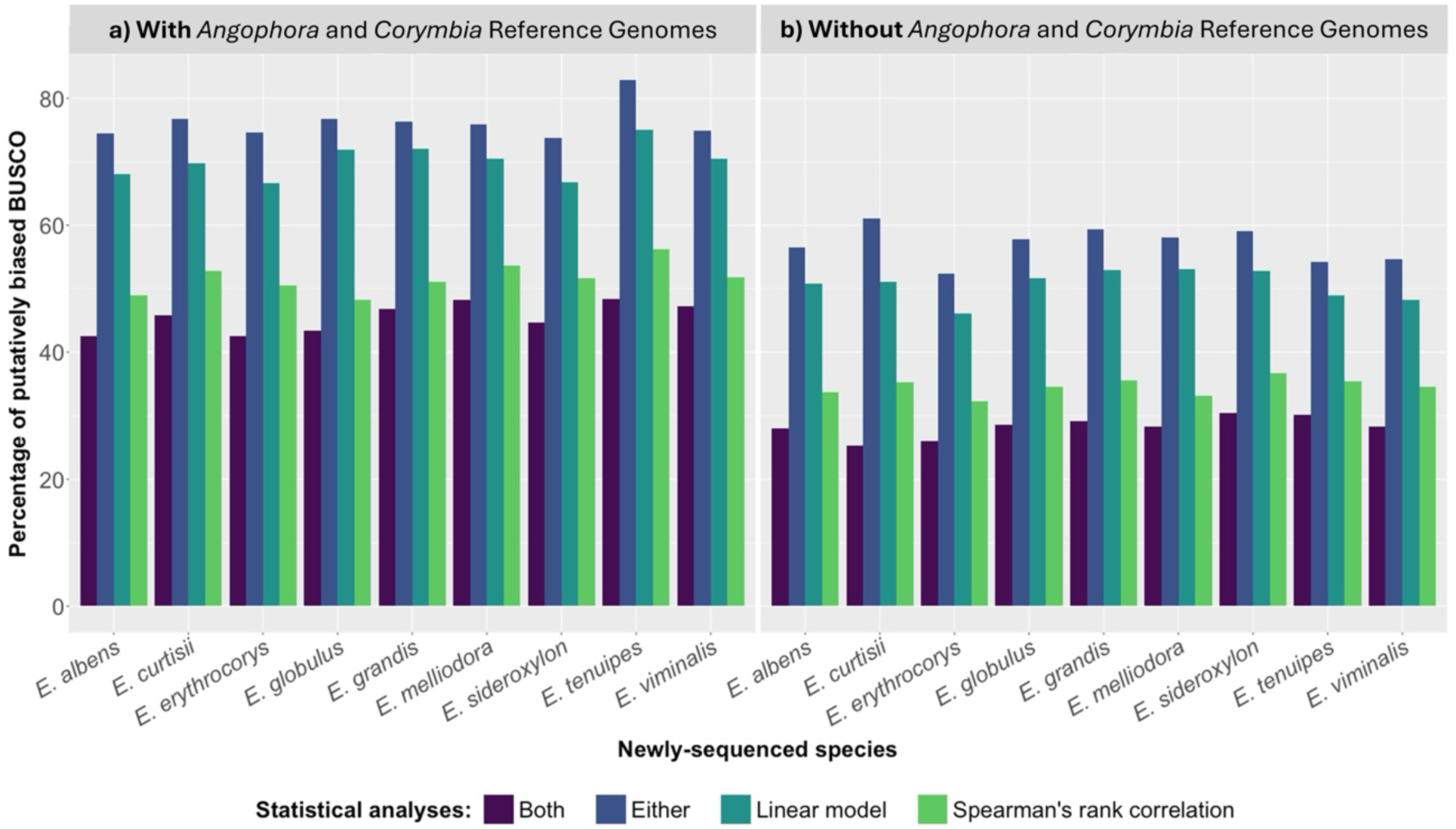
Percentage of putatively biased BUSCO with significant results (*p*<0.05) from linear model and/or Spearman’s rank order correlation test across nine representative *Eucalyptus* species, (a) including or (b) excluding reconstructions mapped to *Angophora* and *Corymbia* reference genomes. Colouring denotes the statistical tests being used.

For example, BUSCO 4838at71240 reconstructed from *E. albens* short-read data shows no detectable reference bias once the *Angophora* / *Corymbia*-based reconstructions are excluded (i.e., non-significant result (*p*≥0.05) based on the linear model), while BUSCO 31772at71240 reconstructed from the same set of short reads shows strong evidence of reference bias (Fig. 3). For BUSCO 4838at71240, all reconstructed BUSCO from *E. albens* (except three that were mapped to *Angophora* and *Corymbia* references) are clustered together with zero phylogenetic distance (Fig. 3a). This is reflected by the results from the linear model; when we excluded the reconstructed BUSCO that were mapped to *Angophora* and *Corymbia* reference genomes, the correlation between the reads and the reference pairwise distances is not statistically significant (R^2^<0.01, *p*=0.182). On the other hand, for BUSCO 31772at71240, some reconstructed loci of *E. albens* appear in the tree very close to the reference genomes that were used to reconstruct them (Fig. 3b). This observation is supported by the linear model, which shows significant correlations between the two distances both when the reconstructed BUSCO that were mapped to *Angophora* and *Corymbia* reference genomes are included (R^2^=0.857, *p*<0.001), and when the *Angophora* and *Corymbia* reference genomes are excluded (R^2^=0.649, *p*<0.001).

**Figure 3.**
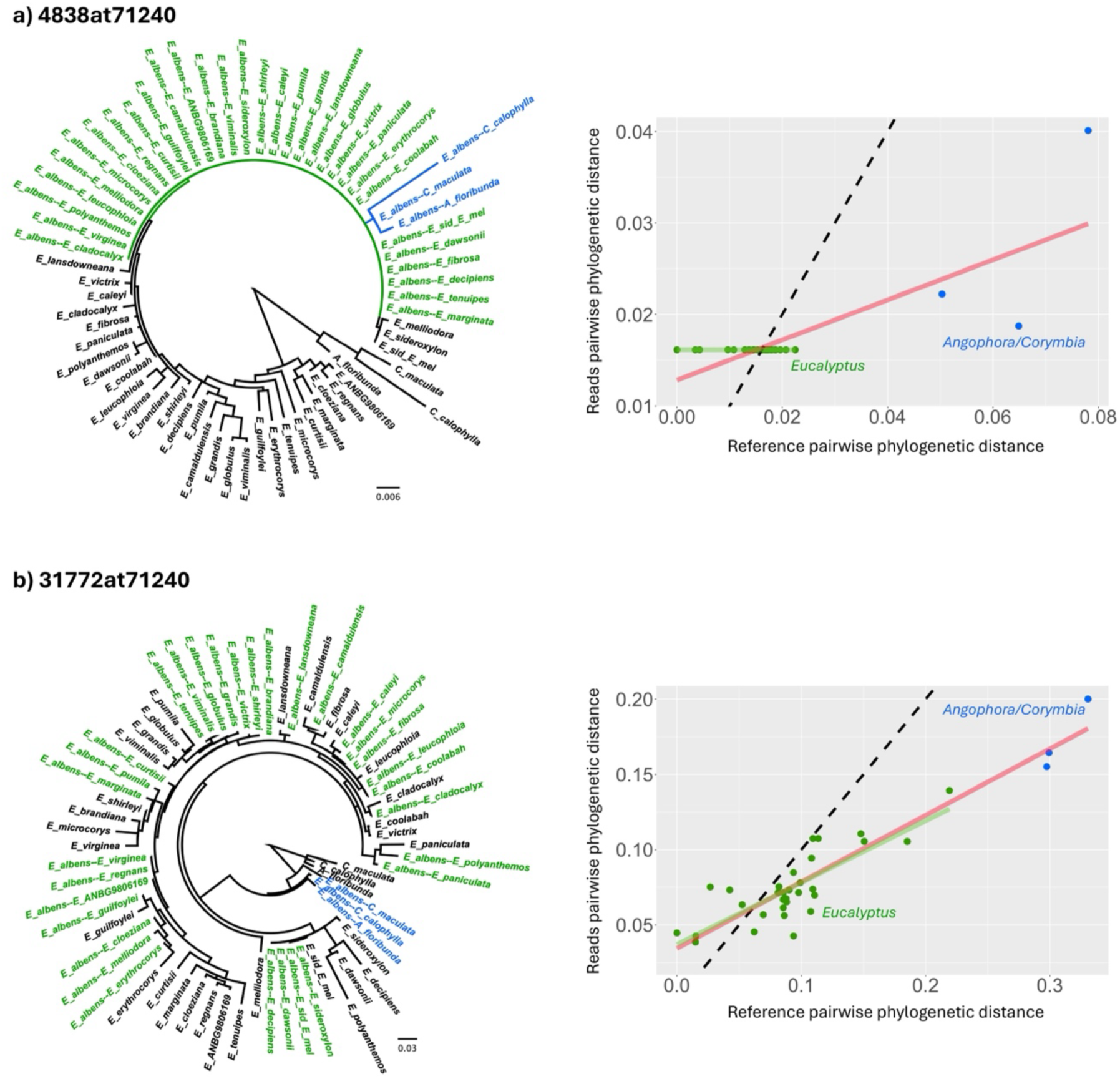
BUSCO (a) 4838at71240 and (b) 31772at71240 for *E. albens* consensus sequences. Left: BUSCO trees; right: scatter plots of the BUSCO pairwise distances to the focal species (i.e., *E. cloeziana*). Dashed line shows x = y line; red solid line shows the fitted line based on a linear model with *Angophora* and *Corymbia* reference genomes; green solid line shows the fitted line based on a linear model without *Angophora* and *Corymbia* reference genomes. Green-coloured tips denote *E. albens* consensus sequences with *Eucalyptus* reference genomes, while the blue-coloured tips denote *E. albens* consensus sequences with either *Corymbia* or *Angophora* reference genomes.

### PhyloRBT flags loci with the most reference bias

To assess the predictive power of PhyloRBT in detecting reference bias, we retrieved the amino acid sequences of individual reconstructed BUSCO and compared them with the sequences inferred from the species’ *de novo* reference genomes. Across the nine representative taxa, there are roughly 4x more sites affected by reference bias than those with other types of error not attributable to reference bias (e.g., heterozygous sites, mapping error, and alignment error; Fig. S2; Table S5), which supports the prevalence of reference bias in this dataset. For each representative species, we then ranked individual BUSCOs based on the proportion of biased sites across reconstructions, and assessed the predictive power of PhyloRBT to detect biased loci. Across the nine representative taxa, considering all loci with a significant result for either the linear model or Spearman’s rank order correlation test had the highest detection power of 80-85% (as it flags the greatest number of putatively biased loci), followed by significance in the linear model alone (74-81%), Spearman’s rank order correlation test alone (58-69%), and finally in requiring significance in both tests (54-65%; Table S6).

To test whether tree-based statistics from PhyloRBT flag the same loci if applied directly to sequences, we repeated the linear model and Spearman’s rank order correlation test on the BUSCO amino acid sequences. We then treated the sequence-based results as the *reference classification* and measured the sensitivity (i.e., the ability to detect true positives) and specificity (i.e., the ability to detect true negatives) of PhyloRBT. We found that the linear model has the strongest agreement between tree-based and sequence-based analyses, having 84% sensitivity and 68% specificity across all BUSCO (Fig. S3). On the other hand, Spearman’s rank order correlation test has a weaker agreement of 58% sensitivity and 65% specificity (Fig. S3). Using *both* tests has similarly low sensitivity of 58% (but 75% specificity), while considering either test has 83% sensitivity and a slightly lower specificity of 57%. Across all loci, PhyloRBT flagged a similar proportion of loci (Table S3) to the sequence-based analysis (65–76% under the linear model and 76–84% under *either* test; Table S7, with *Angophora* + *Corymbia*).

### Excluding putatively biased loci increases the concordance factors of the species clustering

Excluding the putatively biased loci (i.e., BUSCO loci with a significant result from *either* statistical test) generally increases the concordance factors of the reconstruction clustering on the ASTRAL tree compared to when using all BUSCO loci. Specifically, using only the putatively unbiased loci increases the gCF and qCF of the relevant branch by on average 6.11% and 6.08%, respectively, except for *E. globulus* (with 1.04% gCF and 14.25% qCF decrease) and *E. albens* (with 0.57% qCF decrease) (Figs. 4, S4-S12). However, the changes in sCF are more inconsistent, where it increases by 58.05% for *E. tenuipes* when using only putatively unbiased loci, but decreases by 54.15% for *E. curtisii* (Figs. 4, S4-S12).

**Figure 4.**
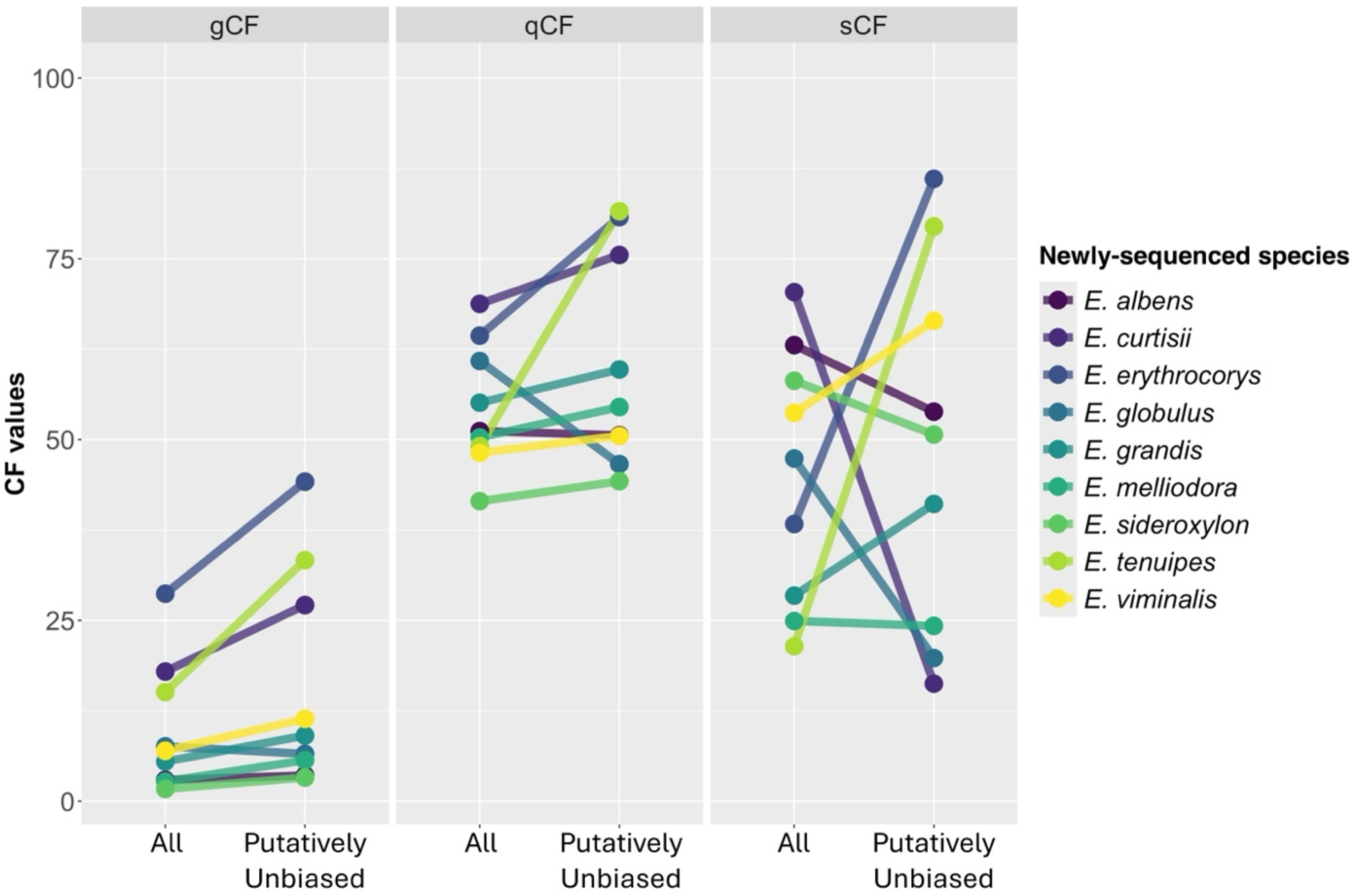
Concordance factors of the relevant branch on the ASTRAL trees inferred using *all* BUSCO loci and only putatively unbiased BUSCO loci. Relevant branch refers to the branch that leads to all reconstructed versions of the species. gCF: gene concordance factors; qCF: quartet concordance factor; sCF: site concordance factor. Colouring denotes different newly-sequenced species.

### Reference bias affects the species placement on the species tree

When only one reconstruction was included in the species tree inference, the placement of the newly-sequenced species varies depending on the reference genome being used. For example, Figures S13-S16 show that the position of *E. albens* reconstruction shifts within the *Adnataria* clade, often with >0.95 posterior probability support even when the placement is inconsistent with the published tree (Ferguson et al. 2024a). This pattern is further supported by calculating the phylogenetic distance between the ASTRAL tree and the published tree. Across the three representative species, Figure 5 (*All*) shows that the newly-sequenced species is placed much deeper in the tree when using reference genomes from *Corymbia* or *Angophora* (except for the short reads of *E. erythrocorys* that is placed at the crown of *Eucalyptus* in the published tree). Conversely, using reference genomes from closely-related taxa pulls the reconstruction closer to the reference genome itself, which recovered the published position for *E. grandis* but not for *E. albens* or *E. erythrocorys* (light green dots in Fig. 5 (*All*)). Excluding the putatively biased BUSCO loci does not consistently reduce the phylogenetic distance to the published tree (Fig. 5 (*Putatively Unbiased*)). However, when we used *both* a reference genome from closely-related taxa during mapping *and* only putatively unbiased BUSCO during the species tree inference, the ASTRAL tree topology is always consistent with the published tree across the three representative species (light green dots in Fig. 5 (*Putatively Unbiased*)).

**Figure 5.**
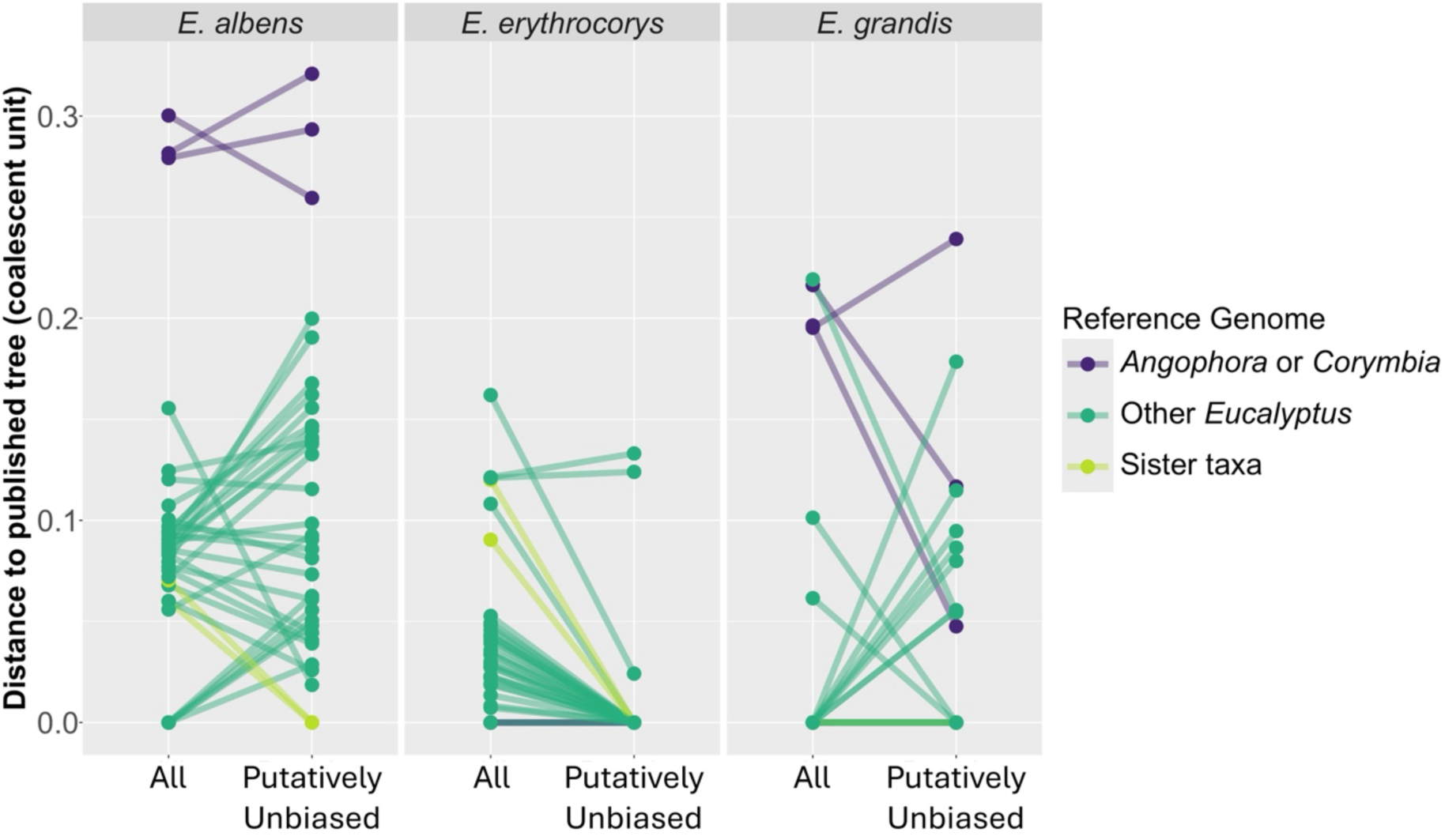
Phylogenetic distance between the position of three *Eucalyptus* species on the ASTRAL trees (inferred using *all* BUSCO loci and only putatively unbiased BUSCO loci) and the published tree (Ferguson et al. 2024a). The y-axis reflects the total branch lengths (in coalescent units) needed for the reconstruction to reach the ‘true’ branch on the published tree (see red-coloured edges on Figs. S13-S16 for reference). Colouring denotes different reference genomes used for mapping (Fig. S1). Lines connect reconstructions with the same reference genome.

To assess if the inconsistent placements of *E. albens* across ASTRAL trees were due to small number of putatively unbiased loci (112 out of 439 BUSCO; Table S3), we repeated the species tree inference but only excluding loci that were statistically significant from *both* statistical tests, which doubled the number of putatively unbiased loci to 252 loci (Table S3). We show that increasing the number of loci reduces the phylogenetic distance to the published tree for most reconstructions (Fig. S17). However, only reconstructions that were mapped to closely-related reference genomes consistently recover the published tree topology (i.e., light green dots in Fig. S17 (*Putatively Unbiased* from *both* tests)). These results further support the use of a closely-related reference genome during mapping, and highlight the importance of addressing reference bias in phylogenomic analyses.

## Discussion

Many phylogenomic studies start by mapping the short-read data of newly-sequenced species to one or more high-quality reference genomes, in order to reconstruct loci for phylogenetic inference. However, reference genome selection has been found to introduce bias that can make the reconstructed locus erroneously similar to the reference genome (Degner et al. 2009; Stevenson et al. 2013; Lin et al. 2024). Using the dataset of Ferguson et al. (2024a), we found that across hundreds of BUSCO loci, more than a quarter showed significant evidence of reference bias across nine *Eucalyptus* species based on PhyloRBT, which was exacerbated when the reference genome became more distantly-related to the newly-sequenced species (Fig. 2; Table S3). This is also consistent when comparing the reconstructed sequences to the sequences inferred from the species’ *de novo* reference genomes (Fig. S2; Table S5). Interestingly, when compared to their closest reference, the proportion of biased sites per 1,000 sites ranges between 3.3 and 5.1 across six representative species (excluding *E. curtisii*, *E. erythrocorys*, and *E. tenuipes* with higher reference distance; Fig. S2). This might represent private alleles (or assembly errors) of the *de novo* reference genomes that are not shared with other reference genomes, and thus are unlikely to be detected by PhyloRBT by design. Crucially, reference bias could affect the position of the newly-sequenced species on the species tree (Figs. S13-S16), which highlights the risk of introducing reference bias in phylogenomic datasets generated using reference mapping, even with closely-related species.

In order to address the bias, the simplest way is to avoid it altogether by assembling the genome *de novo*, which is becoming increasingly affordable with the advancement of long-read sequencing technologies (Ekblom and Wolf 2014; Sedlazeck et al. 2018; Rhie et al. 2021). However, in cases where short sequencing reads are the only data available, *de novo* assembly is particularly challenging (Paszkiewicz and Studholme 2010; Liao et al. 2019), even though previous studies have found success in generating draft genome assemblies from DNA extracted from fresh tissue and museum specimens (e.g., Li et al. 2010; Feigin et al. 2018; Varga et al. 2025). When *de novo* assembly is not feasible and mapping to a reference genome is a necessary part of a phylogenomic study, careful consideration needs to be taken when performing data analyses in the presence of reference bias.

When we used all reconstructions of each of the nine representative *Eucalyptus* species, we found that removing the putatively biased loci from the ASTRAL tree inference generally improved gCF and qCF (but not sCF) of the reconstruction clustering compared to when using *all* BUSCO loci (Fig. 4). In contrast to posterior probability that measures sampling variance, concordance factors (CFs) measure the proportion of informative genes (gCF and qCF) and sites (sCF) that agree with the species tree (Lanfear and Hahn 2024). As a result, posterior probability tends to converge towards 1.0 in phylogenomic datasets due to the large number of loci being used (Gadagkar et al. 2005), while CFs do not monotonically increase with the size of the dataset (Minh et al. 2020a). As PhyloRBT detects reference bias in individual loci, excluding the putatively biased loci reduced the degree of reference bias in the datasets and improved the reconstruction clustering, which was reflected by increased proportion of loci (gCF) and quartets (qCF) supporting the clustering. However, a single locus may contain multiple evolutionary histories due to recombination, such that some sites support the reconstruction clustering while others do not (Degnan and Rosenberg 2009; Mallet et al. 2016; Bryant and Hahn 2020). PhyloRBT does not account for these intra-locus recombination events because we only infer one history (i.e., tree) per locus. Consequently, some sites within the putatively biased loci may in fact be unbiased but were nonetheless excluded from the analysis, and vice versa. As the proportion of sites (sCF) that support the clustering did not always increase after excluding the putatively biased BUSCO loci from the ASTRAL tree inference (Fig. 4), it might suggest that intra-locus recombination is prevalent in our dataset. This is consistent with other data from *Eucalyptus* genomics (Gion et al. 2016; Murray et al. 2019).

Similar to previous findings (Bertels et al. 2014; Valiente-Mullor et al. 2021), we also found that reference bias affected the species tree inference of three representative species (*E. albens*, *E. erythrocorys*, and *E. grandis*), where the placement of the species on the ASTRAL tree was highly dependent on the reference genome being used during mapping (Fig. 5). The usual approach to address reference bias is to map the short reads to a single reference genome from closely-related taxa and use all reconstructed loci to infer the tree. However, we showed that this approach did not consistently recover the published placement of the newly-sequenced species, particularly for *E. albens* and *E. erythrocorys* (i.e., light green dots in the *All* column of Fig. 5). For example, using the reference genomes from *Corymbia* or *Angophora* placed the *E. albens* reconstruction at the crown of *Adnataria* clade with high posterior probability support (Fig. S13). Conversely, when the reference genomes came from closely-related taxa (e.g., *E. melliodora* and *E. sideroxylon*), the *E. albens* reconstruction clustered together with the reference genome itself (Fig. S14). This highlights that reference bias can significantly affect species tree inference, often with high support value, even when the reference genome being used comes from closely-related taxa.

In this study, we flagged a BUSCO locus to be putatively biased if it had a significant result from *either* the linear model or the Spearman’s rank order correlation test, because we wanted to minimise the reference bias contained in the dataset. In addition, this criterion has a relatively high agreement with the sequence-based analyses (Fig. S3). However, considering *either* test also filtered out the biggest number of loci (Table S3), which might increase species tree estimation error due to a reduction in phylogenetic signal. For example, we showed that increasing the number of loci could improve the ASTRAL tree inference (Figs. 5, S17). Therefore, the extent to which putatively biased loci should be removed is likely to depend on the phylogenetic question. If removing every locus flagged by *either* test still leaves sufficient information, it is the preferable choice. However, if this aggressive filtering leaves insufficient information, then users must decide whether the level (and direction) of reference bias in the retained loci is enough to undermine their conclusions. Compared to other tools that assess reference bias of one sample against one reference – such as AMBER through mapping statistics (Dolenz et al. 2024) and `biastools` through departures from the expected allelic balance in a diploid individual (Lin et al. 2024) – PhyloRBT uses the linear model and Spearman’s rank order correlation test to detect phylogenetic *attraction* (i.e., reduced pairwise phylogenetic distance) of individual reconstructions towards their references on individual gene trees. As a result, PhyloRBT is applicable even when no diploid baseline exists, and can be used directly to filter individual loci as long as there is sufficient divergence among samples to infer gene trees. On the other hand, PhyloRBT requires multiple reference genomes (i.e., data points per locus) to make a reliable inference, and cannot analyse each reference individually to identify those with stronger or weaker attraction (unless we repeated the correlation analyses multiple times by excluding one reference at a time). PhyloRBT also benefits from the development of long-read sequencing technologies, as including more reconstructions per locus will increase the reliability of PhyloRBT. However, note that the runtime of PhyloRBT will also increase proportionately with the number of reference genomes used, as each set of short reads needs to be individually mapped to each reference genome.

However, we also showed that when we used the reference genome from closely-related taxa *and* excluded putatively biased BUSCO from the species tree inference, the ASTRAL tree topology became more consistent with the published tree (Ferguson et al. 2024a). Thus, we have two recommendations to address reference bias in phylogenomic datasets when multiple reference genomes are available. First, we should run PhyloRBT to detect loci with strong evidence of reference bias and exclude them from the species tree inference. Second, for each of the remaining loci, we should select the reconstruction that was mapped to the closest reference genome (which can be estimated directly from the short sequencing reads using tools such as Mash (Ondov et al. 2016) in the absence of a published species tree). Alternatively, we could reconstruct the remaining loci directly from the short reads of the newly-sequenced species using HybPiper (Johnson et al. 2016) or other similar tools that allow for iterative mapping during locus reconstructions (Hahn et al. 2013). In brief, iterative mapping involves: (i) mapping short reads to the reference locus, (ii) constructing a new consensus sequence from the mapped reads, and (iii) using this consensus sequence as the updated target locus. By repeating these steps until convergence (i.e., until no new reads are mapped), the reconstructed locus should be less affected by reference bias and more accurately reflect the sequence of the newly-sequenced species.

Overall, our study shows one example of the prevalence of reference bias in phylogenomic datasets and thus proposes PhyloRBT to detect the bias on individual loci using multiple reference genomes. We achieved this by individually mapping each set of short reads to multiple reference genomes, reconstructing the locus of interest from the consensus sequences, and then running correlation analyses between the reads pairwise distance (i.e., the pairwise phylogenetic distance between the consensus sequence and the focal species) and the reference pairwise distance (i.e., the pairwise phylogenetic distance between the reference genome and the focal species). While we only presented one use case of our method – namely, to infer species tree using BUSCO loci – it can also be used for other phylogenomic analyses (e.g., inference of gene trees), or even extended to evaluate different approaches that aim to mitigate reference bias during upstream analyses (e.g., selection of variant calling parameters (Shafer et al. 2017; Barbitoff et al. 2022; Rick et al. 2024), realignment of reads (Homer and Nelson 2010; Pirooznia et al. 2014; Mose et al. 2019), iterative mapping (Dutilh et al. 2009; Tsai et al. 2010; Hahn et al. 2013), and the use of pan-genomes (The Computational Pan-Genomics Consortium 2018; Valenzuela et al. 2018; Chen et al. 2021)). Moreover, the pattern we observed -where using more distantly-related references would increase the proportion of putatively biased loci from PhyloRBT -is also applicable for other groups of taxa, since the degree of reference bias should correlate with the genetic divergence between the newly-sequenced sample and the reference being used (Gopalakrishnan et al. 2017; Bohling 2020; Huang et al. 2024). This pattern also extends to the intra-specific level (Degner et al. 2009; Günther and Nettelblad 2019; Thorburn et al. 2023). However, as PhyloRBT relies on gene trees to detect reference bias, it is likely to have limited power in intraspecific datasets since divergence between conspecific samples is likely to be low, at least at the level of individual loci. In such cases, tools such as AMBER (Dolenz et al. 2024) and biastools (Lin et al. 2024) might be better suited. In conclusion, we believe that PhyloRBT provides a valuable resource to detect reference bias in empirical datasets using multiple reference genomes, which can help to reduce its impact on downstream analyses, particularly phylogenetic inference.

## Data Availability

Summary tables and datasets reported in this study will be available upon the acceptance of the paper.

## Supplementary Materials

Supplementary material will be available online upon the acceptance of the paper.

## Acknowledgements

We thank Minh Bui, Justin Borevitz, the members of the Lanfear and Bui Lab from Australian National University, as well as Kevin Murray from the Max Planck Institute for comments on the study design and the manuscript.

