## Supplementary Materials for "PhyloRBT: A Phylogenetic Approach to Detect Reference Bias in Phylogenomic Datasets"

**Supplementary Tables**

**Table S1.** Mean mapping coverage for each pair of representative species (as columns) and reference genome (as rows).

| reference | *E. albens* | *E. curtisii* | *E. erythrocorys* | *E. globulus* | *E. grandis* | *E. melliodora* | *E. sideroxylon* | *E. tenuipes* | *E. viminalis* |
| --- | --- | --- | --- | --- | --- | --- | --- | --- | --- |
| *A. floribunda* | 14.375 | 9.081 | 11.898 | 13.205 | 13.231 | 10.288 | 19.795 | 8.535 | 9.990 |
| *C. calophylla* | 14.688 | 9.221 | 12.189 | 13.469 | 13.488 | 10.592 | 20.298 | 8.739 | 10.203 |
| *C. maculata* | 13.828 | 8.748 | 11.463 | 12.663 | 12.700 | 9.887 | 19.082 | 8.214 | 9.585 |
| *E. ANBG9806169* | 21.836 | 12.806 | 17.800 | 21.343 | 20.853 | 16.557 | 29.819 | 12.393 | 16.881 |
| *E. albens* | N/A | 10.885 | 15.466 | 19.710 | 19.200 | 16.045 | 29.044 | 10.490 | 15.834 |
| *E. brandiana* | 24.248 | 12.706 | 18.047 | 23.379 | 22.622 | 18.150 | 33.023 | 12.263 | 18.734 |
| *E. caleyi* | 21.471 | 11.019 | 15.713 | 20.315 | 19.692 | 16.035 | 29.219 | 10.634 | 16.334 |
| *E. camaldulensis* | 21.673 | 11.495 | 16.410 | 21.948 | 21.239 | 16.294 | 29.573 | 11.101 | 17.638 |
| *E. cladocalyx* | 22.533 | 11.729 | 16.785 | 21.795 | 21.073 | 16.823 | 30.625 | 11.344 | 17.500 |
| *E. cloeziana* | 23.247 | 13.629 | 18.933 | 22.669 | 22.145 | 17.572 | 31.761 | 13.241 | 17.930 |
| *E. coolabah* | 20.832 | 10.765 | 15.283 | 19.692 | 19.150 | 15.717 | 28.592 | 10.398 | 15.791 |
| *E. curtisii* | 26.040 | N/A | 21.140 | 25.185 | 24.635 | 20.406 | 36.247 | 15.031 | 19.947 |
| *E. dawsonii* | 18.373 | 9.612 | 13.523 | 17.025 | 16.660 | 14.455 | 25.605 | 9.219 | 13.712 |
| *E. decipiens* | 20.887 | 11.152 | 15.831 | 20.206 | 19.665 | 15.938 | 28.669 | 10.753 | 16.216 |
| *E. erythrocorys* | 19.850 | 11.595 | N/A | 19.364 | 18.838 | 15.054 | 27.206 | 11.283 | 15.251 |
| *E. fibrosa* | 21.663 | 11.278 | 15.839 | 20.311 | 19.840 | 16.589 | 29.870 | 10.832 | 16.277 |
| *E. globulus* | 22.123 | 11.774 | 16.811 | N/A | 21.752 | 16.640 | 30.203 | 11.375 | 18.341 |
| *E. grandis* | 20.087 | 10.689 | 15.239 | 20.141 | N/A | 15.265 | 27.543 | 10.326 | 16.228 |
| *E. guilfoylei* | 24.744 | 14.240 | 19.802 | 24.098 | 23.569 | 19.385 | 34.387 | 13.708 | 19.223 |
| *E. lansdowneana* | 20.492 | 10.625 | 14.938 | 18.978 | 18.527 | 15.854 | 28.321 | 10.201 | 15.258 |
| *E. leucophloia* | 22.102 | 11.652 | 16.373 | 20.934 | 20.425 | 17.045 | 30.597 | 11.196 | 16.765 |
| *E. marginata* | 21.524 | 12.546 | 17.556 | 21.260 | 20.667 | 16.135 | 29.251 | 12.184 | 16.856 |
| *E. melliodora* | 20.246 | 10.493 | 14.817 | 18.849 | 18.376 | N/A | 28.053 | 10.063 | 15.209 |
| *E. microcorys* | 26.193 | 14.792 | 20.814 | 25.748 | 25.014 | 19.862 | 35.823 | 14.280 | 20.531 |
| *E. paniculata* | 21.629 | 11.281 | 15.969 | 20.351 | 19.797 | 16.552 | 29.772 | 10.840 | 16.340 |
| *E. polyanthemos* | 21.405 | 11.097 | 15.659 | 19.853 | 19.409 | 16.502 | 29.619 | 10.660 | 15.960 |
| *E. pumila* | 23.301 | 12.614 | 17.702 | 23.107 | 22.475 | 18.276 | 32.441 | 12.086 | 18.628 |
| *E. regnans* | 22.183 | 13.032 | 18.102 | 21.721 | 21.184 | 16.794 | 30.308 | 12.639 | 17.196 |
| *E. shirleyi* | 21.264 | 11.117 | 15.745 | 20.093 | 19.566 | 16.344 | 29.339 | 10.682 | 16.160 |
| *E. sideroxylon x E. melliodora* | 20.926 | 10.805 | 15.378 | 19.854 | 19.268 | 15.769 | 28.923 | 10.425 | 15.919 |
| *E. sideroxylon* | 21.217 | 10.846 | 15.491 | 20.048 | 19.435 | 15.874 | N/A | 10.469 | 16.065 |
| *E. tenuipes* | 27.473 | 16.516 | 22.658 | 26.985 | 26.221 | 20.809 | 37.564 | N/A | 21.426 |
| *E. victrix* | 22.517 | 11.689 | 16.564 | 21.328 | 20.709 | 17.034 | 30.847 | 11.244 | 17.137 |
| *E. viminalis* | 21.821 | 11.644 | 16.574 | 22.329 | 21.359 | 16.586 | 29.839 | 11.206 | N/A |
| *E. virginea* | 22.934 | 12.256 | 17.474 | 22.420 | 21.765 | 17.284 | 31.297 | 11.830 | 17.972 |

**Table S2.** Percentage of mapped reads for each pair of representative species (as columns) and reference genome (as rows).

| reference | *E. albens* | *E. curtisii* | *E. erythrocorys* | *E. globulus* | *E. grandis* | *E. melliodora* | *E. sideroxylon* | *E. tenuipes* | *E. viminalis* |
| --- | --- | --- | --- | --- | --- | --- | --- | --- | --- |
| *A. floribunda* | 56.89 | 60.14 | 52.68 | 55.72 | 56.99 | 42.18 | 58.29 | 60.66 | 48.62 |
| *C. calophylla* | 58.82 | 61.84 | 54.49 | 57.30 | 58.54 | 44.22 | 60.30 | 62.49 | 49.97 |
| *C. maculata* | 57.18 | 60.31 | 52.94 | 55.72 | 57.03 | 42.45 | 58.73 | 60.87 | 48.53 |
| *E. ANBG9806169* | 89.81 | 92.71 | 86.41 | 92.42 | 92.28 | 85.68 | 91.82 | 93.43 | 88.06 |
| *E. albens* | N/A | 93.23 | 88.52 | 96.71 | 96.11 | 94.63 | 97.55 | 94.08 | 93.70 |
| *E. brandiana* | 95.12 | 92.65 | 87.25 | 96.21 | 95.42 | 92.19 | 96.21 | 93.40 | 92.90 |
| *E. caleyi* | 95.96 | 93.01 | 87.77 | 96.71 | 96.01 | 93.72 | 97.03 | 93.71 | 93.64 |
| *E. camaldulensis* | 93.28 | 92.62 | 87.80 | 97.44 | 96.62 | 91.62 | 95.91 | 93.35 | 94.46 |
| *E. cladocalyx* | 95.15 | 92.53 | 87.35 | 96.28 | 95.58 | 92.17 | 96.22 | 93.35 | 93.02 |
| *E. cloeziana* | 90.18 | 93.34 | 86.74 | 92.72 | 92.54 | 86.53 | 92.47 | 94.27 | 88.25 |
| *E. coolabah* | 96.21 | 93.24 | 87.98 | 96.71 | 96.13 | 94.39 | 97.49 | 94.11 | 93.59 |
| *E. curtisii* | 89.89 | N/A | 86.22 | 92.72 | 92.13 | 86.23 | 92.15 | 94.07 | 88.45 |
| *E. dawsonii* | 96.58 | 93.60 | 88.87 | 96.79 | 96.24 | 95.11 | 97.79 | 94.34 | 93.76 |
| *E. decipiens* | 95.29 | 93.18 | 88.42 | 96.57 | 95.94 | 92.82 | 96.48 | 94.04 | 93.42 |
| *E. erythrocorys* | 88.19 | 90.74 | N/A | 90.66 | 90.12 | 83.09 | 90.23 | 92.03 | 85.92 |
| *E. fibrosa* | 96.10 | 93.19 | 87.93 | 96.82 | 96.20 | 94.00 | 97.15 | 93.90 | 93.69 |
| *E. globulus* | 93.12 | 92.48 | 87.72 | N/A | 96.53 | 91.45 | 95.75 | 93.27 | 94.93 |
| *E. grandis* | 94.20 | 93.47 | 88.79 | 97.85 | N/A | 93.38 | 96.86 | 94.26 | 95.11 |
| *E. guilfoylei* | 92.12 | 93.21 | 87.26 | 94.53 | 93.95 | 88.67 | 93.69 | 93.94 | 90.91 |
| *E. lansdowneana* | 96.36 | 93.41 | 88.35 | 96.85 | 96.17 | 94.66 | 97.55 | 94.22 | 93.70 |
| *E. leucophloia* | 95.44 | 92.90 | 87.79 | 96.53 | 95.86 | 93.06 | 96.57 | 93.67 | 93.28 |
| *E. marginata* | 90.10 | 92.73 | 86.42 | 92.70 | 92.31 | 86.01 | 92.13 | 93.59 | 88.51 |
| *E. melliodora* | 96.26 | 93.19 | 88.28 | 96.78 | 96.14 | N/A | 97.53 | 93.92 | 93.69 |
| *E. microcorys* | 92.24 | 92.51 | 86.94 | 94.24 | 93.53 | 87.87 | 93.25 | 93.44 | 90.51 |
| *E. paniculata* | 95.87 | 93.05 | 88.23 | 96.68 | 95.93 | 93.43 | 96.89 | 93.82 | 93.47 |
| *E. polyanthemos* | 96.48 | 93.49 | 88.52 | 96.80 | 96.26 | 94.95 | 97.75 | 94.22 | 93.79 |
| *E. pumila* | 94.84 | 92.91 | 88.00 | 97.25 | 96.42 | 92.16 | 96.04 | 93.63 | 94.39 |
| *E. regnans* | 89.24 | 92.46 | 85.93 | 92.04 | 91.77 | 85.67 | 91.81 | 93.37 | 87.72 |
| *E. shirleyi* | 95.73 | 93.15 | 88.42 | 96.78 | 96.10 | 93.78 | 97.05 | 93.88 | 93.66 |
| *E. sideroxylon x E. melliodora* | 94.49 | 92.96 | 88.02 | 96.73 | 96.01 | 93.67 | 97.31 | 93.71 | 93.62 |
| *E. sideroxylon* | 95.59 | 92.41 | 87.76 | 96.37 | 95.64 | 93.27 | N/A | 93.21 | 93.19 |
| *E. tenuipes* | 88.47 | 92.97 | 85.70 | 91.51 | 91.07 | 83.76 | 90.48 | N/A | 86.97 |
| *E. victrix* | 95.58 | 92.70 | 87.70 | 96.47 | 95.71 | 93.11 | 96.65 | 93.38 | 93.31 |
| *E. viminalis* | 94.89 | 92.68 | 87.87 | 97.90 | 96.74 | 91.91 | 95.94 | 93.41 | N/A |
| *E. virginea* | 94.94 | 93.07 | 88.28 | 96.59 | 95.90 | 92.05 | 96.07 | 93.86 | 93.22 |

**Table S3.** Proportion of BUSCO with significant statistical test results (*p*<0.05) across representative taxa. lm: linear model; sp: Spearman’s rank correlation. Values are percentages of the total number of BUSCO (#).

| Species | # | With *Angophora* + *Corymbia* | | | | Without *Angophora* + *Corymbia* | | | |
| --- | --- | --- | --- | --- | --- | --- | --- | --- | --- |
|  |  | lm | sp | both | either | lm | sp | both | either |
| *E. albens* | 439 | 68.11 | 48.97 | 42.60 | 74.49 | 50.80 | 33.71 | 28.02 | 56.49 |
| *E. curtisii* | 301 | 69.77 | 52.82 | 45.85 | 76.74 | 51.16 | 35.22 | 25.25 | 61.13 |
| *E. erythrocorys* | 338 | 66.57 | 50.59 | 42.60 | 74.56 | 46.15 | 32.25 | 26.04 | 52.37 |
| *E. globulus* | 263 | 71.86 | 48.29 | 43.35 | 76.81 | 51.71 | 34.60 | 28.52 | 57.79 |
| *E. grandis* | 419 | 72.08 | 51.07 | 46.78 | 76.37 | 52.98 | 35.56 | 29.12 | 59.43 |
| *E. melliodora* | 365 | 70.41 | 53.70 | 48.22 | 75.89 | 53.15 | 33.15 | 28.22 | 58.08 |
| *E. sideroxylon* | 466 | 66.74 | 51.72 | 44.64 | 73.82 | 52.79 | 36.70 | 30.47 | 59.01 |
| *E. tenuipes* | 192 | 75.00 | 56.25 | 48.44 | 82.81 | 48.96 | 35.42 | 30.21 | 54.17 |
| *E. viminalis* | 315 | 70.48 | 51.75 | 47.30 | 74.92 | 48.25 | 34.60 | 28.25 | 54.60 |

**Table S4.** Proportion of BUSCO trimmed using trimAl with significant statistical test results (*p*<0.05) across representative taxa. lm: linear model; sp: Spearman’s rank correlation. Values are percentages of the total number of BUSCO (#).

| Species | # | With *Angophora* + *Corymbia* | | | | Without *Angophora* + *Corymbia* | | | |
| --- | --- | --- | --- | --- | --- | --- | --- | --- | --- |
|  |  | lm | sp | both | either | lm | sp | both | either |
| *E. albens* | 439 | 67.65 | 48.75 | 43.74 | 72.67 | 51.25 | 31.44 | 27.79 | 54.90 |
| *E. curtisii* | 301 | 67.77 | 56.81 | 49.17 | 75.42 | 49.50 | 31.89 | 24.25 | 57.14 |
| *E. erythrocorys* | 338 | 65.09 | 50.59 | 43.79 | 71.89 | 47.34 | 32.84 | 27.22 | 52.96 |
| *E. globulus* | 263 | 69.96 | 52.09 | 45.63 | 76.43 | 50.57 | 34.22 | 28.14 | 56.65 |
| *E. grandis* | 419 | 72.08 | 54.42 | 49.88 | 76.61 | 50.60 | 33.65 | 28.40 | 55.85 |
| *E. melliodora* | 365 | 67.67 | 52.60 | 47.12 | 73.15 | 53.15 | 35.62 | 30.96 | 57.81 |
| *E. sideroxylon* | 466 | 65.02 | 51.29 | 45.71 | 70.60 | 51.29 | 33.69 | 26.61 | 58.37 |
| *E. tenuipes* | 192 | 70.31 | 58.85 | 48.44 | 80.73 | 40.63 | 35.42 | 26.04 | 50.00 |
| *E. viminalis* | 315 | 67.30 | 54.60 | 47.30 | 74.60 | 48.57 | 34.92 | 28.25 | 55.24 |

**Table S5.** Number of sites that differ from the *de novo* reference sequences across all reconstructed BUSCO for each representative species. #: number of unique BUSCO per species; n_sites: total number of sites; n_valid: total number of sites, excluding gaps and unknown amino acids; n_divergent: number of sites that differ between the *de novo* reference and the reference used for mapping; n_biased: number of *n_divergent* sites where the reconstruction follows the reference it was mapped to; n_other: number of *n_valid* sites where the reconstruction does not follow either reference.

| Species | # | n_sites | n_valid | n_divergent | n_biased | n_other |
| --- | --- | --- | --- | --- | --- | --- |
| *E. albens* | 425 | 9046584 | 7880460 | 237462 | 55017 | 15237 |
| *E. curtisii* | 293 | 6327468 | 5498798 | 202376 | 38661 | 10584 |
| *E. erythrocorys* | 337 | 6567746 | 5753347 | 267011 | 39904 | 10541 |
| *E. globulus* | 261 | 4933196 | 4309001 | 158824 | 30682 | 7000 |
| *E. grandis* | 404 | 8451244 | 7408385 | 246139 | 45120 | 13143 |
| *E. melliodora* | 344 | 7823876 | 6905006 | 191134 | 38281 | 10233 |
| *E. sideroxylon* | 456 | 9696800 | 8453970 | 245922 | 52817 | 15177 |
| *E. tenuipes* | 191 | 3457222 | 2963155 | 127304 | 26620 | 6720 |
| *E. viminalis* | 309 | 7003354 | 6103599 | 208268 | 39873 | 11833 |

### **Table S6.** Predictive power of PhyloRBT based on the top-*N* most-biased loci. #: number of BUSCO; significant test: statistical test with *p*<0.05 (lm: linear model; sp: Spearman’s rank correlation test); *N*: number of putatively biased loci flagged by PhyloRBT; flagged: *N* as a percentage of #; caught: percentage of *flagged* BUSCO in the top-*N* most-biased loci.

| Species | # | significant test | *N* | flagged (%) | caught (%) |
| --- | --- | --- | --- | --- | --- |
| *E. albens* | 425 | lm | 287 | 68 | 77 |
|  |  | sp | 206 | 48 | 58 |
|  |  | both | 179 | 42 | 54 |
|  |  | either | 314 | 74 | 82 |
| *E. curtisii* | 293 | lm | 204 | 70 | 74 |
|  |  | sp | 156 | 53 | 61 |
|  |  | both | 135 | 46 | 56 |
|  |  | either | 225 | 77 | 80 |
| *E. erythrocorys* | 337 | lm | 225 | 67 | 76 |
|  |  | sp | 171 | 51 | 59 |
|  |  | both | 144 | 43 | 58 |
|  |  | either | 252 | 75 | 81 |
| *E. globulus* | 261 | lm | 189 | 72 | 80 |
|  |  | sp | 127 | 49 | 60 |
|  |  | both | 114 | 44 | 58 |
|  |  | either | 202 | 77 | 84 |
| *E. grandis* | 404 | lm | 291 | 72 | 80 |
|  |  | sp | 207 | 51 | 62 |
|  |  | both | 189 | 47 | 60 |
|  |  | either | 309 | 76 | 83 |
| *E. melliodora* | 344 | lm | 239 | 69 | 77 |
|  |  | sp | 182 | 53 | 65 |
|  |  | both | 162 | 47 | 59 |
|  |  | either | 259 | 75 | 82 |
| *E. sideroxylon* | 456 | lm | 303 | 66 | 78 |
|  |  | sp | 235 | 52 | 60 |
|  |  | both | 202 | 44 | 58 |
|  |  | either | 336 | 74 | 82 |
| *E. tenuipes* | 191 | lm | 143 | 75 | 81 |
|  |  | sp | 108 | 57 | 65 |
|  |  | both | 93 | 49 | 58 |
|  |  | either | 158 | 83 | 85 |
| *E. viminalis* | 309 | lm | 219 | 71 | 77 |
|  |  | sp | 161 | 52 | 69 |
|  |  | both | 147 | 48 | 65 |
|  |  | either | 233 | 75 | 82 |

**Table S7.** Proportion of BUSCO with significant statistical test results (*p*<0.05) across representative taxa when applying PhyloRBT’s tests on the amino acid p-distances. lm: linear model; sp: Spearman’s rank correlation. Values are percentages of the total number of BUSCO (#).

| Species | # | With *Angophora* + *Corymbia* | | | | Without *Angophora* + *Corymbia* | | | |
| --- | --- | --- | --- | --- | --- | --- | --- | --- | --- |
|  |  | lm | sp | both | either | lm | sp | both | either |
| *E. albens* | 425 | 71.76 | 70.82 | 61.88 | 80.71 | 52.24 | 48.24 | 40.47 | 60.00 |
| *E. curtisii* | 293 | 75.43 | 71.33 | 64.16 | 82.59 | 59.73 | 49.49 | 41.30 | 67.92 |
| *E. erythrocorys* | 337 | 65.28 | 62.61 | 51.63 | 76.26 | 51.04 | 40.65 | 32.64 | 59.05 |
| *E. globulus* | 261 | 72.03 | 73.18 | 62.45 | 82.76 | 54.79 | 44.83 | 38.70 | 60.92 |
| *E. grandis* | 404 | 73.51 | 66.58 | 59.90 | 80.20 | 54.46 | 47.03 | 41.58 | 59.90 |
| *E. melliodora* | 344 | 75.87 | 73.26 | 65.41 | 83.72 | 56.10 | 51.74 | 43.02 | 64.83 |
| *E. sideroxylon* | 456 | 68.42 | 71.05 | 59.65 | 79.82 | 52.19 | 50.44 | 39.69 | 62.94 |
| *E. tenuipes* | 191 | 71.20 | 70.68 | 59.69 | 82.20 | 54.97 | 47.64 | 39.27 | 63.35 |
| *E. viminalis* | 309 | 71.20 | 75.40 | 63.11 | 83.50 | 57.61 | 53.72 | 43.04 | 68.28 |

### **Supplementary Figures**

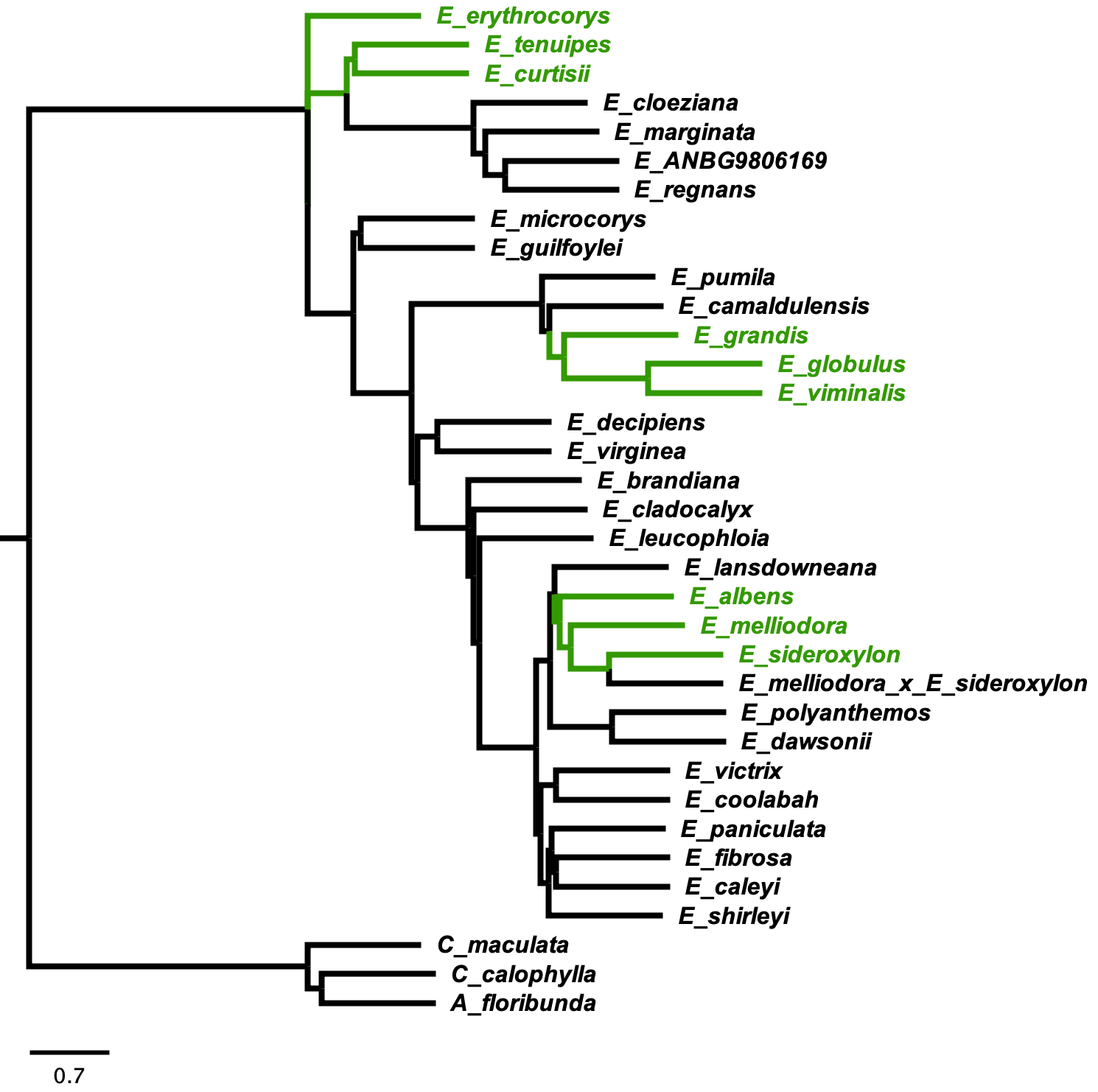

**Figure S1.** Species tree of the 35 eucalypt species from Ferguson et al. (2024a). Tree is rooted on *Corymbia* and *Angophora*. Coloured tips represent the nine representative *Eucalyptus* taxa that are the focus of this study.

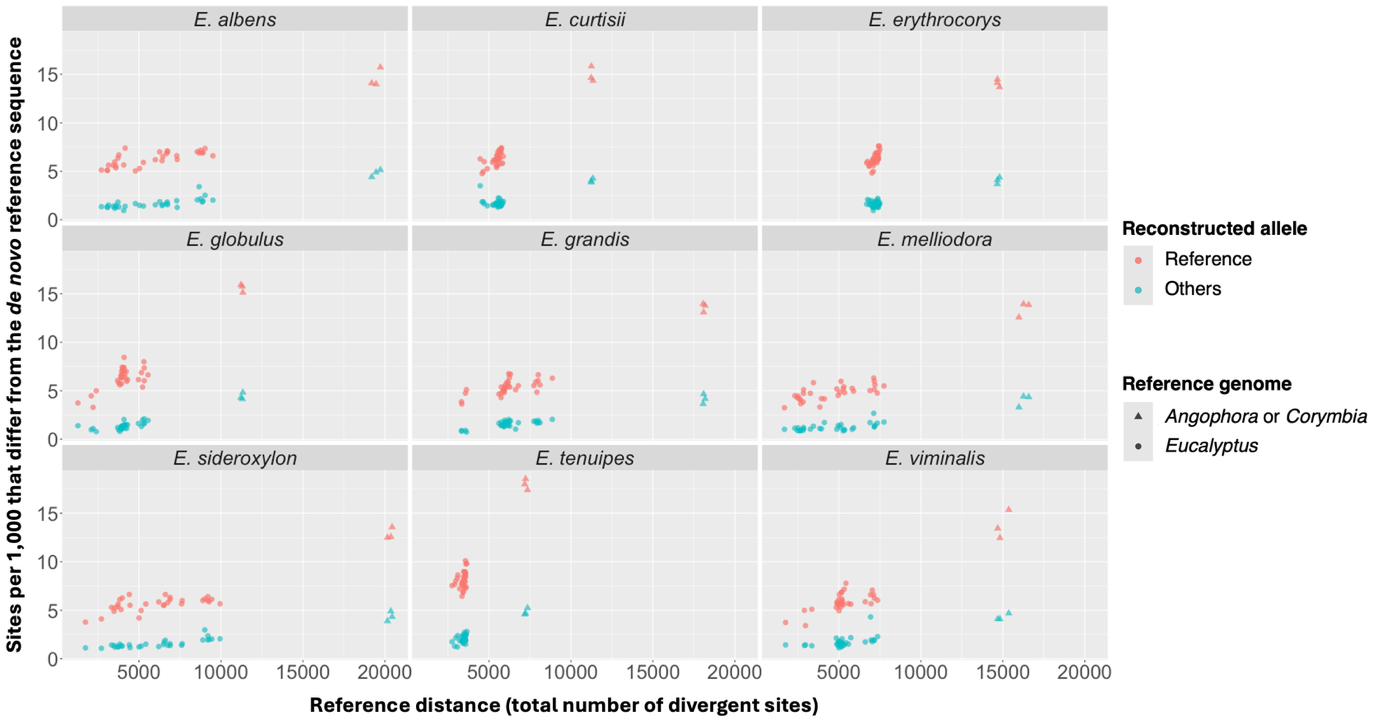

**Figure S2.** Correlation between the total number of divergent sites (i.e., reference distance) and the total number of sites (per 1,000) that differ from the *de novo* reference sequence across all BUSCO. Colours denote alleles carried by the reconstructions, while shapes denote different reference genomes.

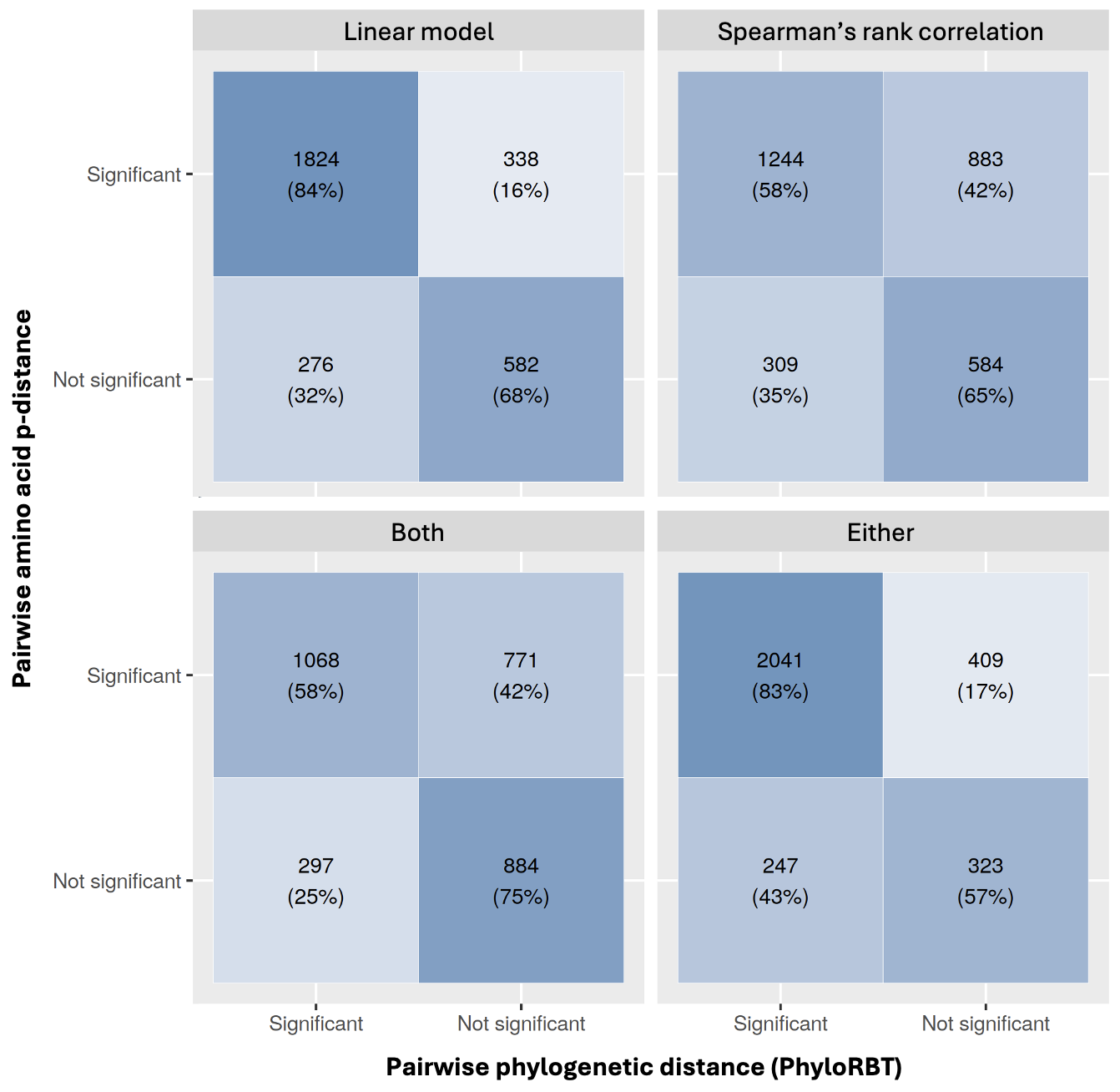

**Figure S3.** Confusion matrices of putatively biased BUSCO based on sequence-based (i.e., amino acid p-distance) and tree-based correlation analyses (i.e., phylogenetic distance from PhyloRBT) across the nine representative *Eucalyptus* taxa. Each matrix represents statistical test(s) used. The top number on each square denotes the total number of BUSCO across nine representative taxa, while the bottom number denotes its percentage.

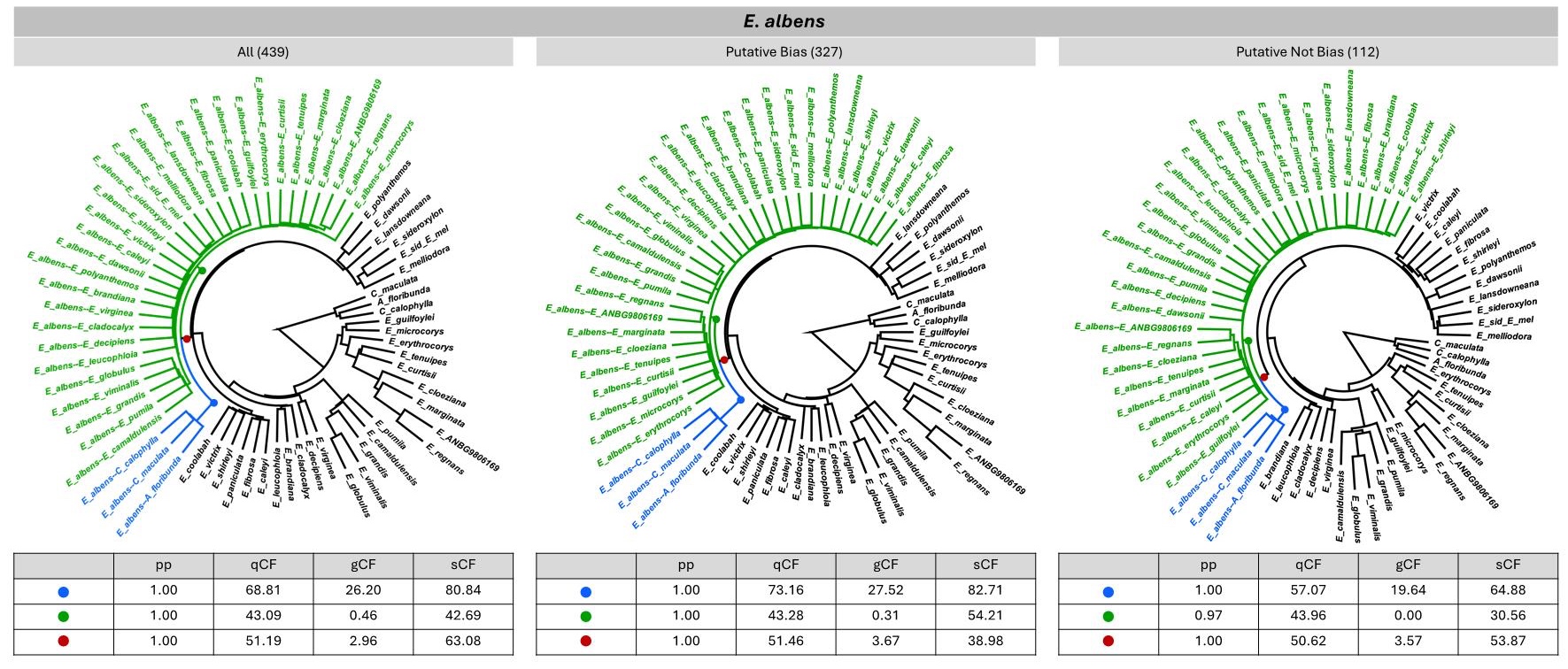

**Figure S4.** ASTRAL species trees and branch supports for the short-read data of *Eucalyptus albens* using three different sets of BUSCO loci: all BUSCO; BUSCO with significant results from either or both statistical tests; and BUSCO with non-significant results from both statistical tests. Numbers in parentheses show the number of BUSCO. Green colour denotes consensus sequences with *Eucalyptus* reference genomes; blue colour denotes consensus sequences with *Angophora* or *Corymbia* reference genomes; red colour denotes the most recent common ancestor of *E. albens* consensus sequences.

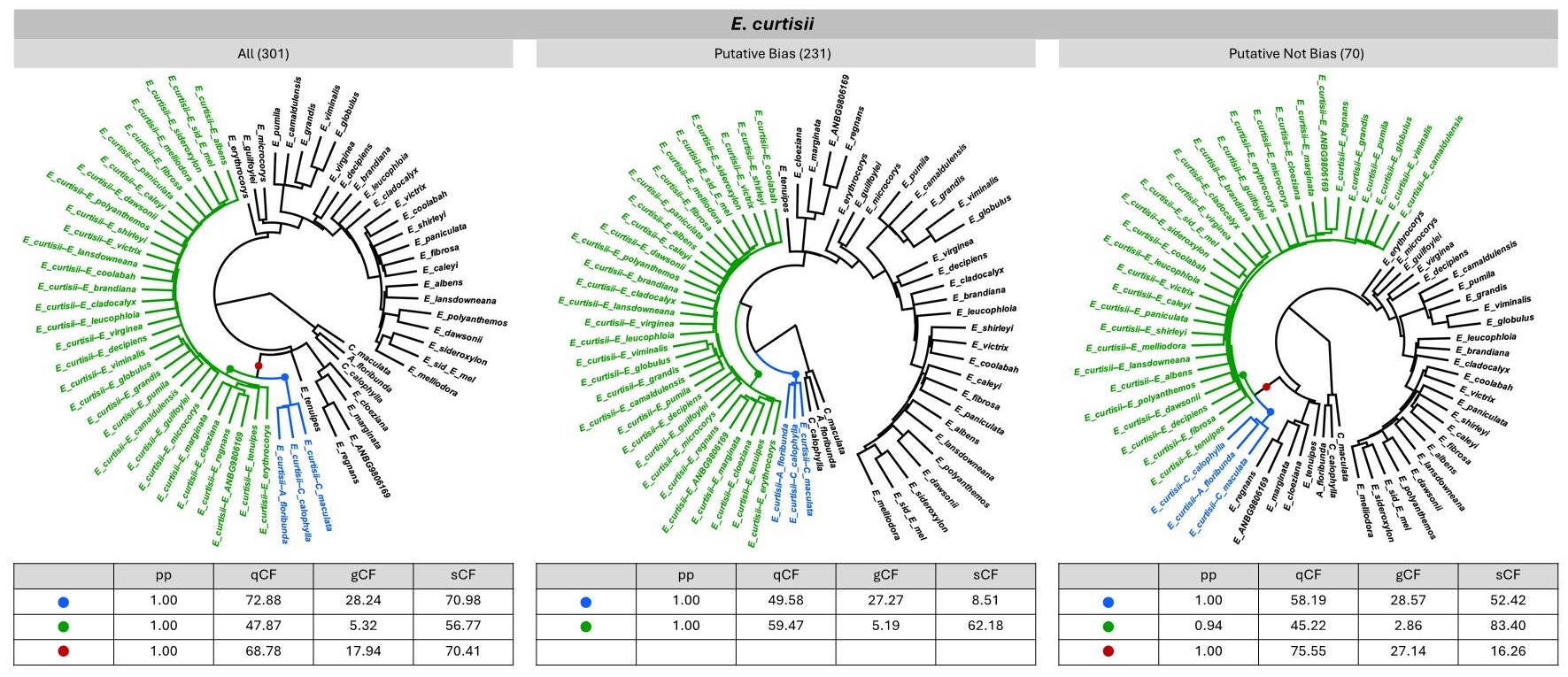

**Figure S5.** ASTRAL species trees and branch supports for the short-read data of *Eucalyptus curtisii* using three different sets of BUSCO loci: all BUSCO; BUSCO with significant results from either or both statistical tests; and BUSCO with non-significant results from both statistical tests. Numbers in parentheses show the number of BUSCO. Green colour denotes consensus sequences with *Eucalyptus* reference genomes; blue colour denotes consensus sequences with *Angophora* or *Corymbia* reference genomes; red colour denotes the most recent common ancestor of *E. curtisii* consensus sequences.

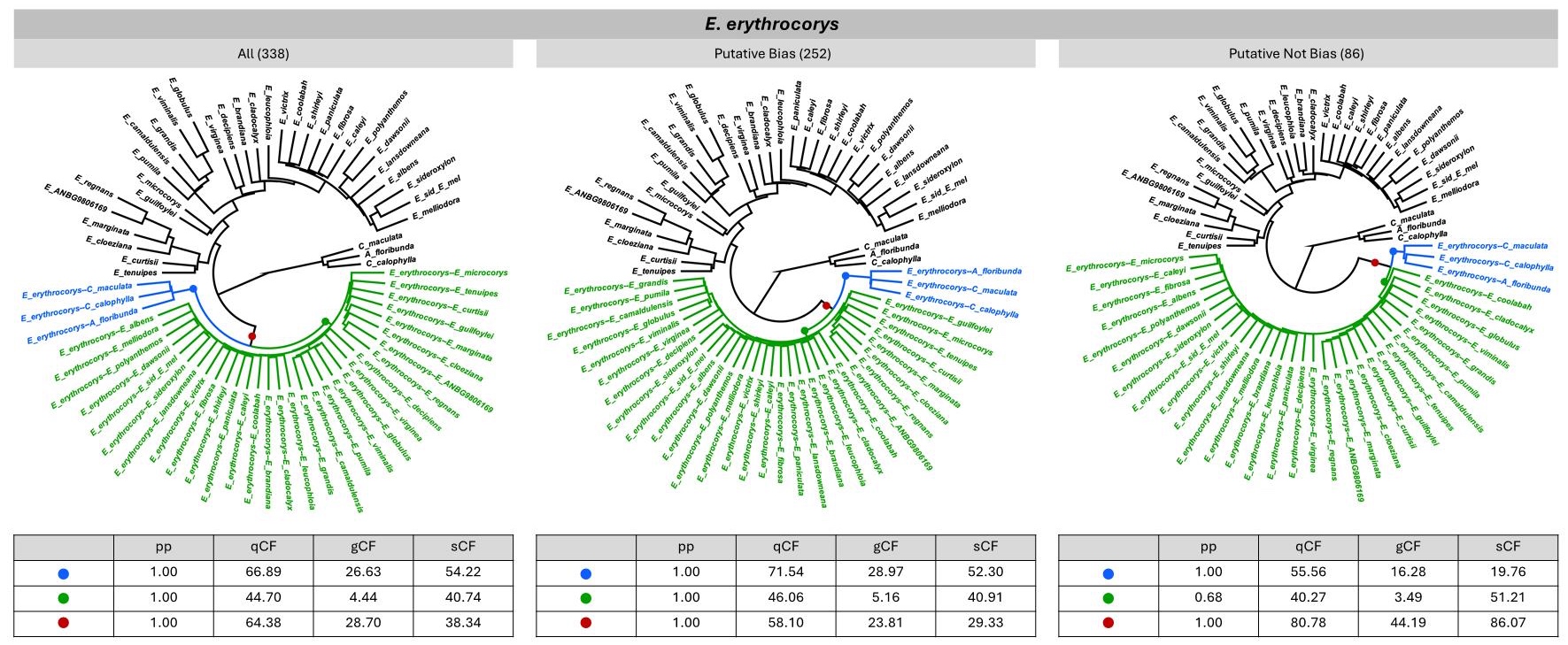

**Figure S6.** ASTRAL species trees and branch supports for the short-read data of *Eucalyptus erythrocorys* using three different sets of BUSCO loci: all BUSCO; BUSCO with significant results from either or both statistical tests; and BUSCO with non-significant results from both statistical tests. Numbers in parentheses show the number of BUSCO. Green colour denotes consensus sequences with *Eucalyptus* reference genomes; blue colour denotes consensus sequences with *Angophora* or *Corymbia* reference genomes; red colour denotes the most recent common ancestor of *E. erythrocorys* consensus sequences.

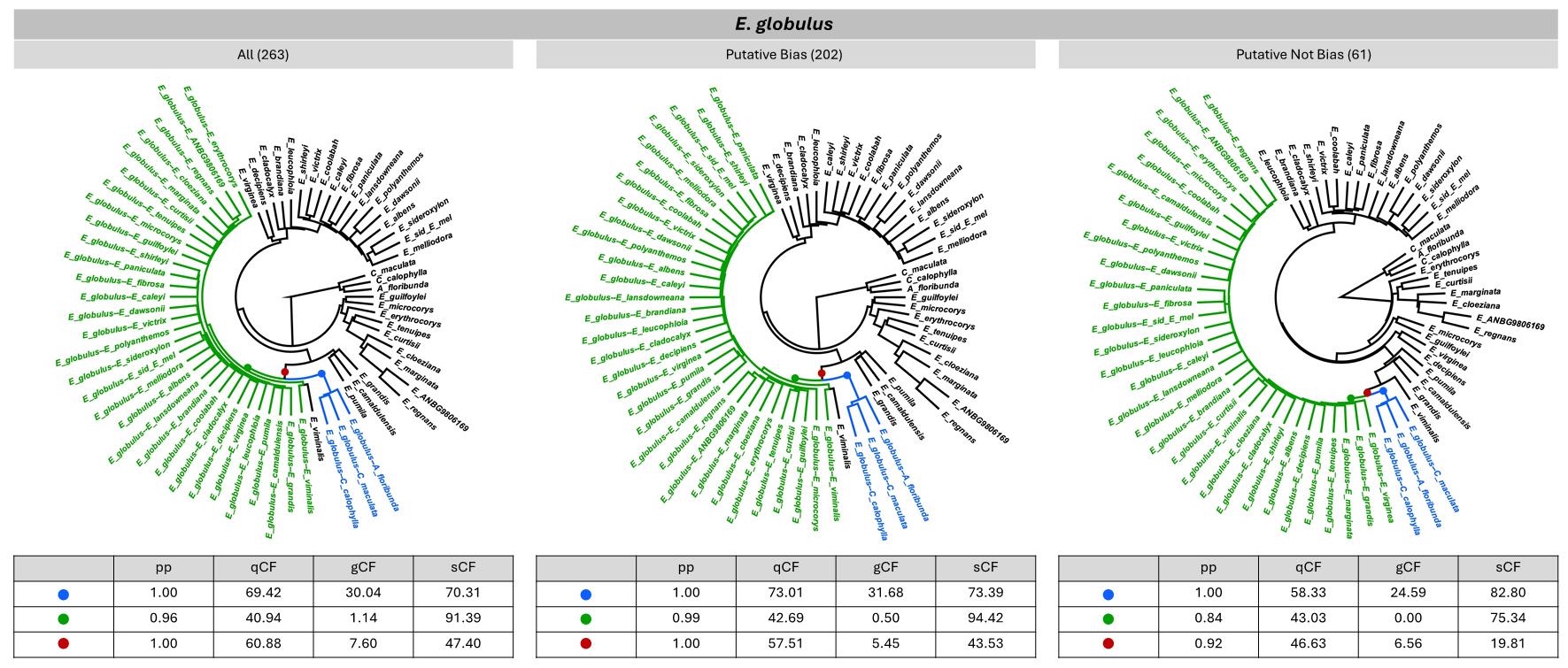

**Figure S7.** ASTRAL species trees and branch supports for the short-read data of *Eucalyptus globulus* using three different sets of BUSCO loci: all BUSCO; BUSCO with significant results from either or both statistical tests; and BUSCO with non-significant results from both statistical tests. Numbers in parentheses show the number of BUSCO. Green colour denotes consensus sequences with *Eucalyptus* reference genomes; blue colour denotes consensus sequences with *Angophora* or *Corymbia* reference genomes; red colour denotes the most recent common ancestor of *E. globulus* consensus sequences.

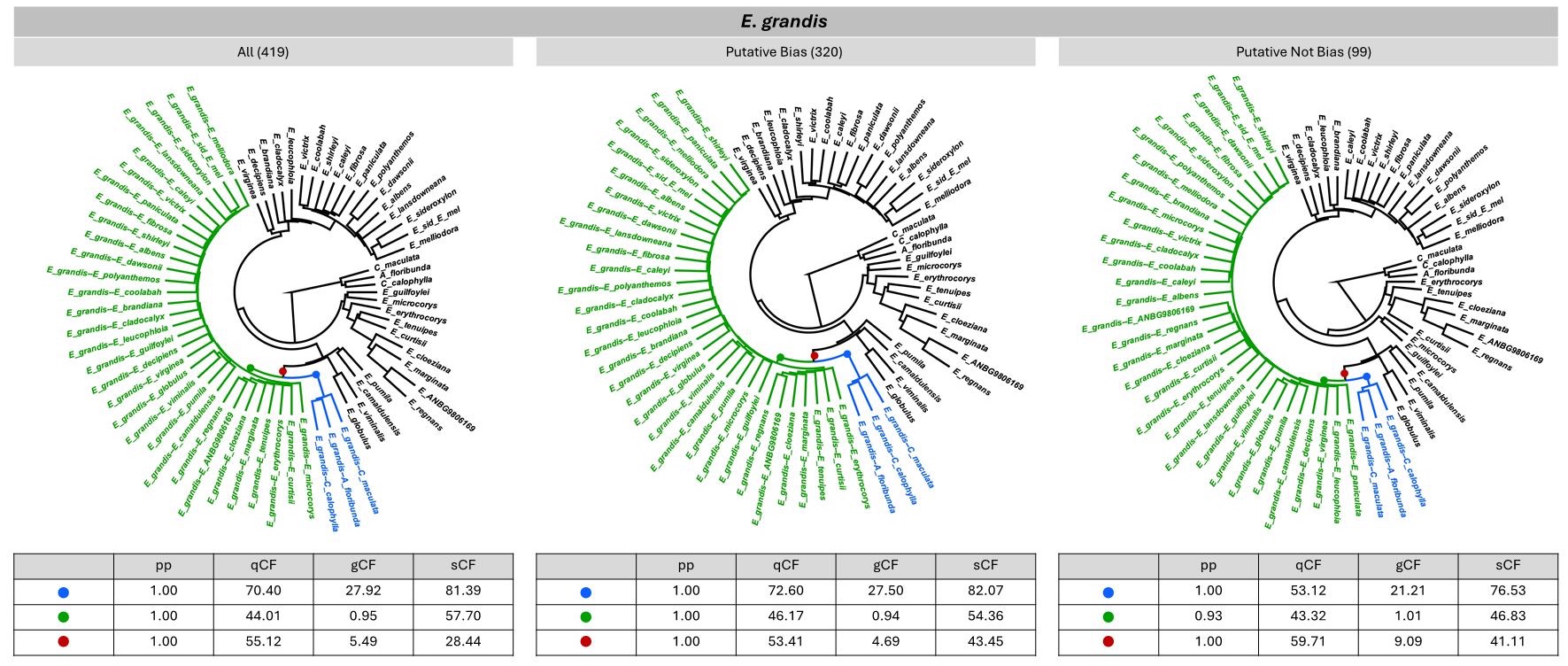

**Figure S8.** ASTRAL species trees and branch supports for the short-read data of *Eucalyptus grandis* using three different sets of BUSCO loci: all BUSCO; BUSCO with significant results from either or both statistical tests; and BUSCO with non-significant results from both statistical tests. Numbers in parentheses show the number of BUSCO. Green colour denotes consensus sequences with *Eucalyptus* reference genomes; blue colour denotes consensus sequences with *Angophora* or *Corymbia* reference genomes; red colour denotes the most recent common ancestor of *E. grandis* consensus sequences.

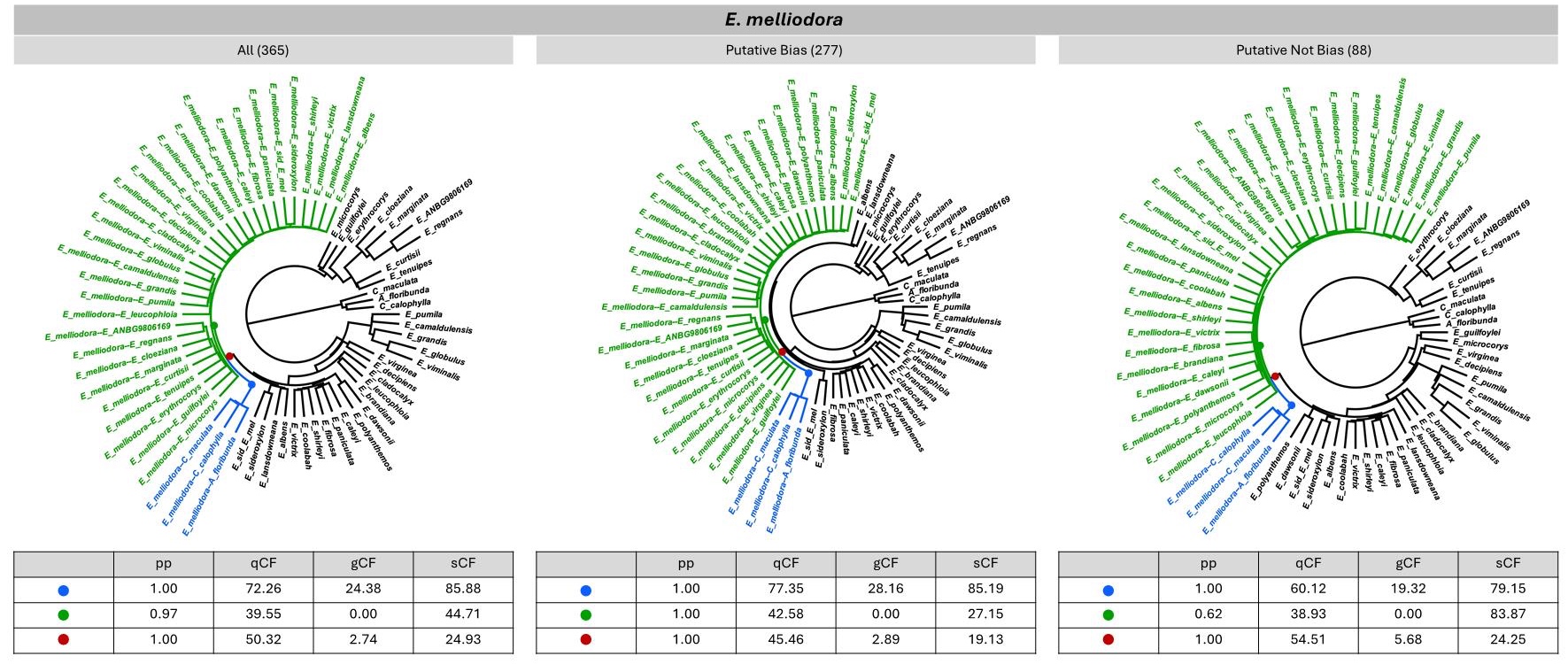

**Figure S9.** ASTRAL species trees and branch supports for the short-read data of *Eucalyptus melliodora* using three different sets of BUSCO loci: all BUSCO; BUSCO with significant results from either or both statistical tests; and BUSCO with non-significant results from both statistical tests. Numbers in parentheses show the number of BUSCO. Green colour denotes consensus sequences with *Eucalyptus* reference genomes; blue colour denotes consensus sequences with *Angophora* or *Corymbia* reference genomes; red colour denotes the most recent common ancestor of *E. melliodora* consensus sequences.

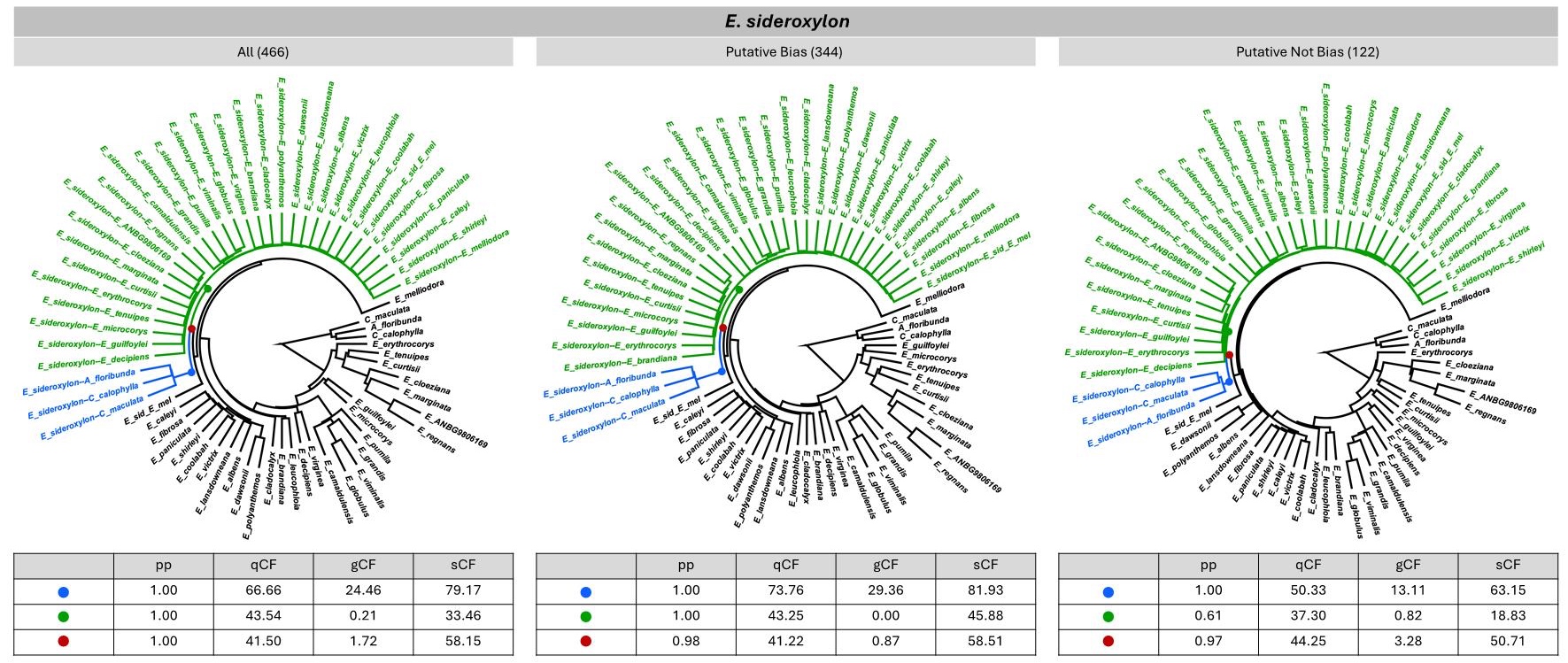

**Figure S10.** ASTRAL species trees and branch supports for the short-read data of *Eucalyptus sideroxylon* using three different sets of BUSCO loci: all BUSCO; BUSCO with significant results from either or both statistical tests; and BUSCO with non-significant results from both statistical tests. Numbers in parentheses show the number of BUSCO. Green colour denotes consensus sequences with *Eucalyptus* reference genomes; blue colour denotes consensus sequences with *Angophora* or *Corymbia* reference genomes; red colour denotes the most recent common ancestor of *E. sideroxylon* consensus sequences.

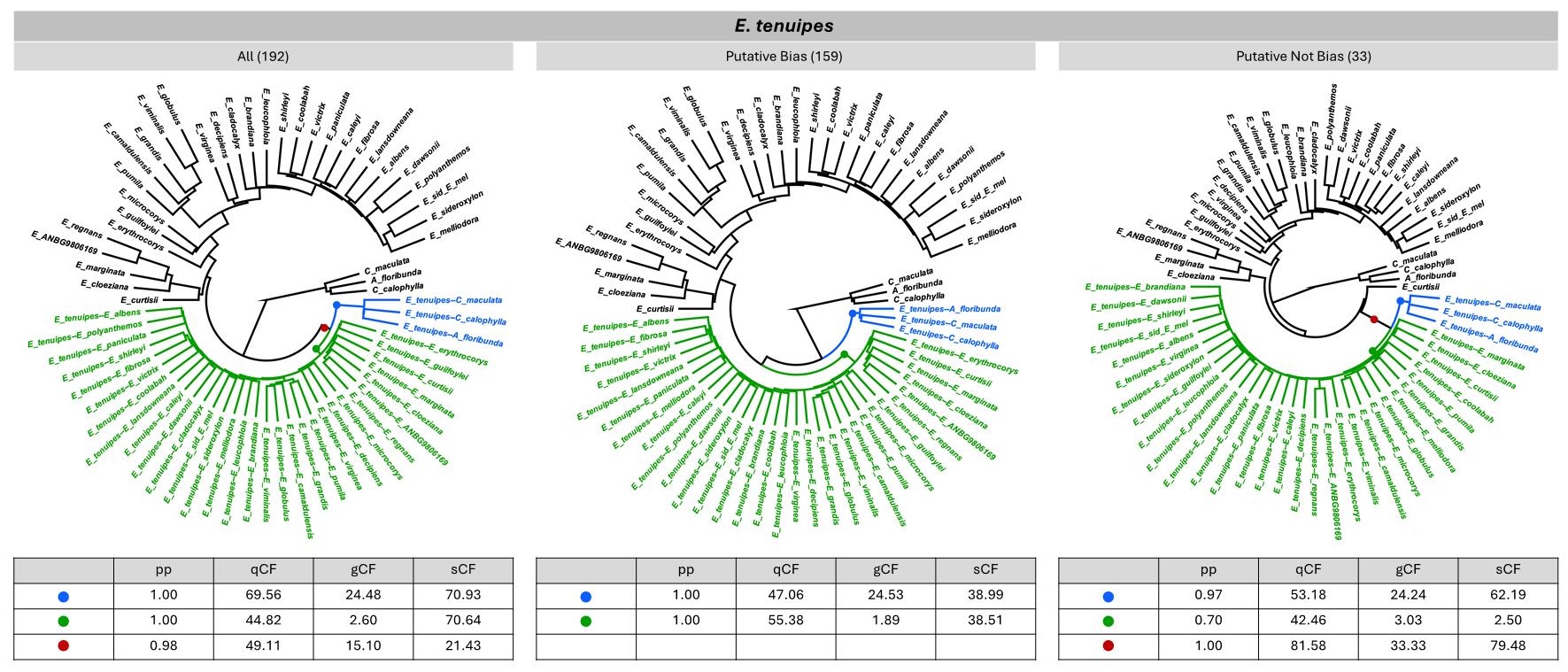

**Figure S11.** ASTRAL species trees and branch supports for the short-read data of *Eucalyptus tenuipes* using three different sets of BUSCO loci: all BUSCO; BUSCO with significant results from either or both statistical tests; and BUSCO with non-significant results from both statistical tests. Numbers in parentheses show the number of BUSCO. Green colour denotes consensus sequences with *Eucalyptus* reference genomes; blue colour denotes consensus sequences with *Angophora* or *Corymbia* reference genomes; red colour denotes the most recent common ancestor of *E. tenuipes* consensus sequences.

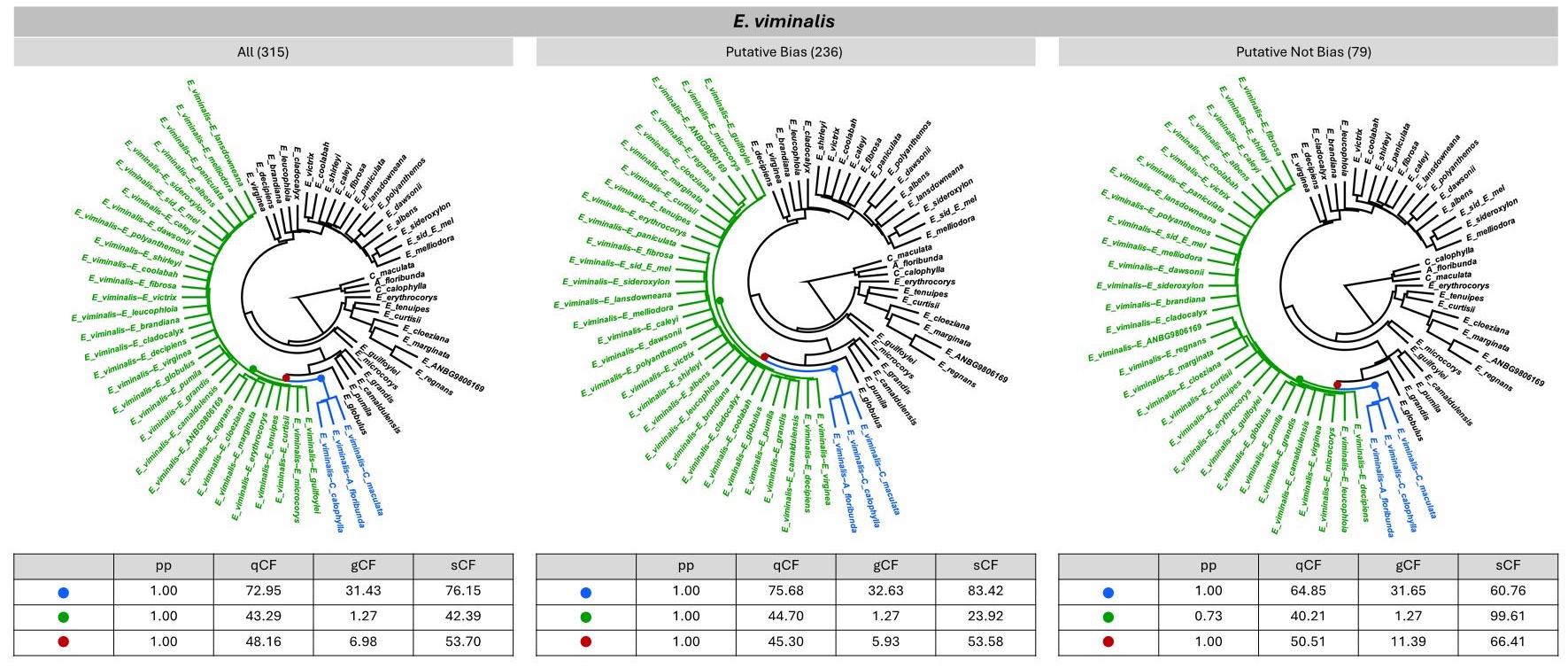

**Figure S12.** ASTRAL species trees and branch supports for the short-read data of *Eucalyptus viminalis* using three different sets of BUSCO loci: all BUSCO; BUSCO with significant results from either or both statistical tests; and BUSCO with non-significant results from both statistical tests. Numbers in parentheses show the number of BUSCO. Green colour denotes consensus sequences with *Eucalyptus* reference genomes; blue colour denotes consensus sequences with *Angophora* or *Corymbia* reference genomes; red colour denotes the most recent common ancestor of *E. viminalis* consensus sequences.

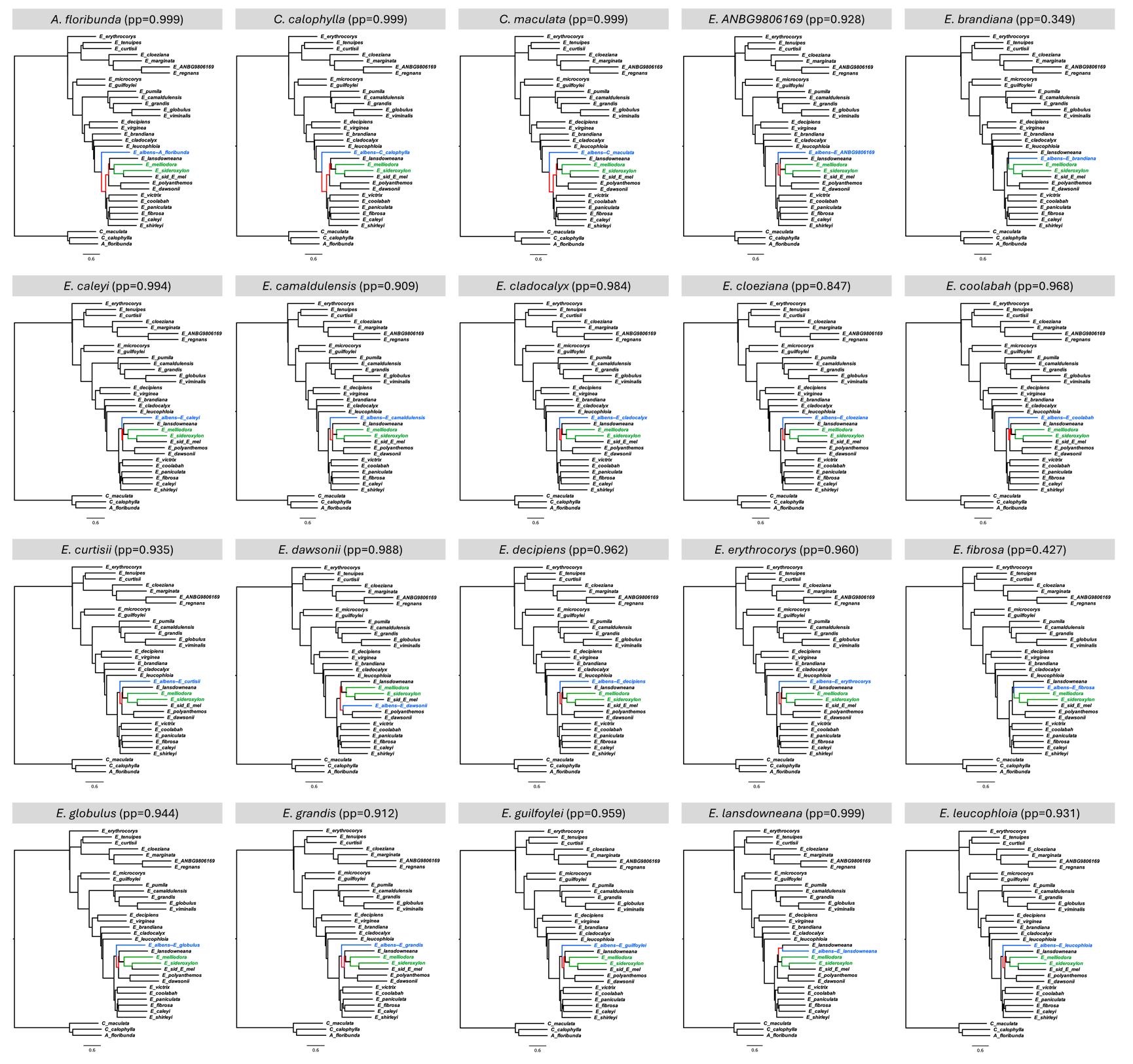

**Figure S13.** ASTRAL species trees for the short-read data of *Eucalyptus albens* across different reference genomes based on all 439 BUSCO (part 1). pp: ASTRAL posterior probability. Blue colour denotes *E. albens*; green colour denotes sister taxa of *E. albens* according to the published tree (Fig. S1); red colour denotes the positional shift of *E. albens* compared to the published tree (Fig. S1).

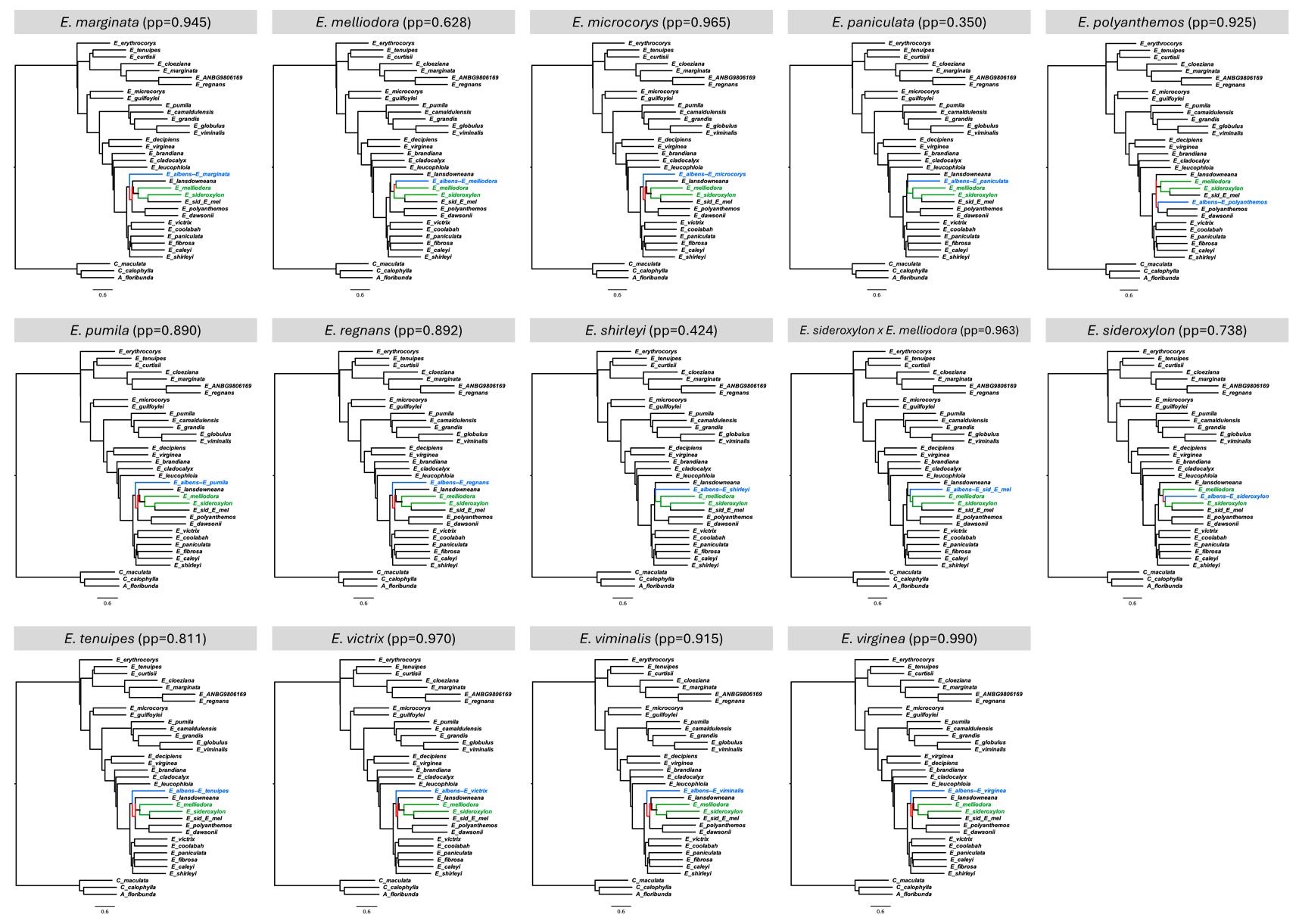

**Figure S14.** ASTRAL species trees for the short-read data of *Eucalyptus albens* across different reference genomes based on all 439 BUSCO (part 2). pp: ASTRAL posterior probability. Blue colour denotes *E. albens*; green colour denotes sister taxa of *E. albens* according to the published tree (Fig. S1); red colour denotes the positional shift of *E. albens* compared to the published tree (Fig. S1).

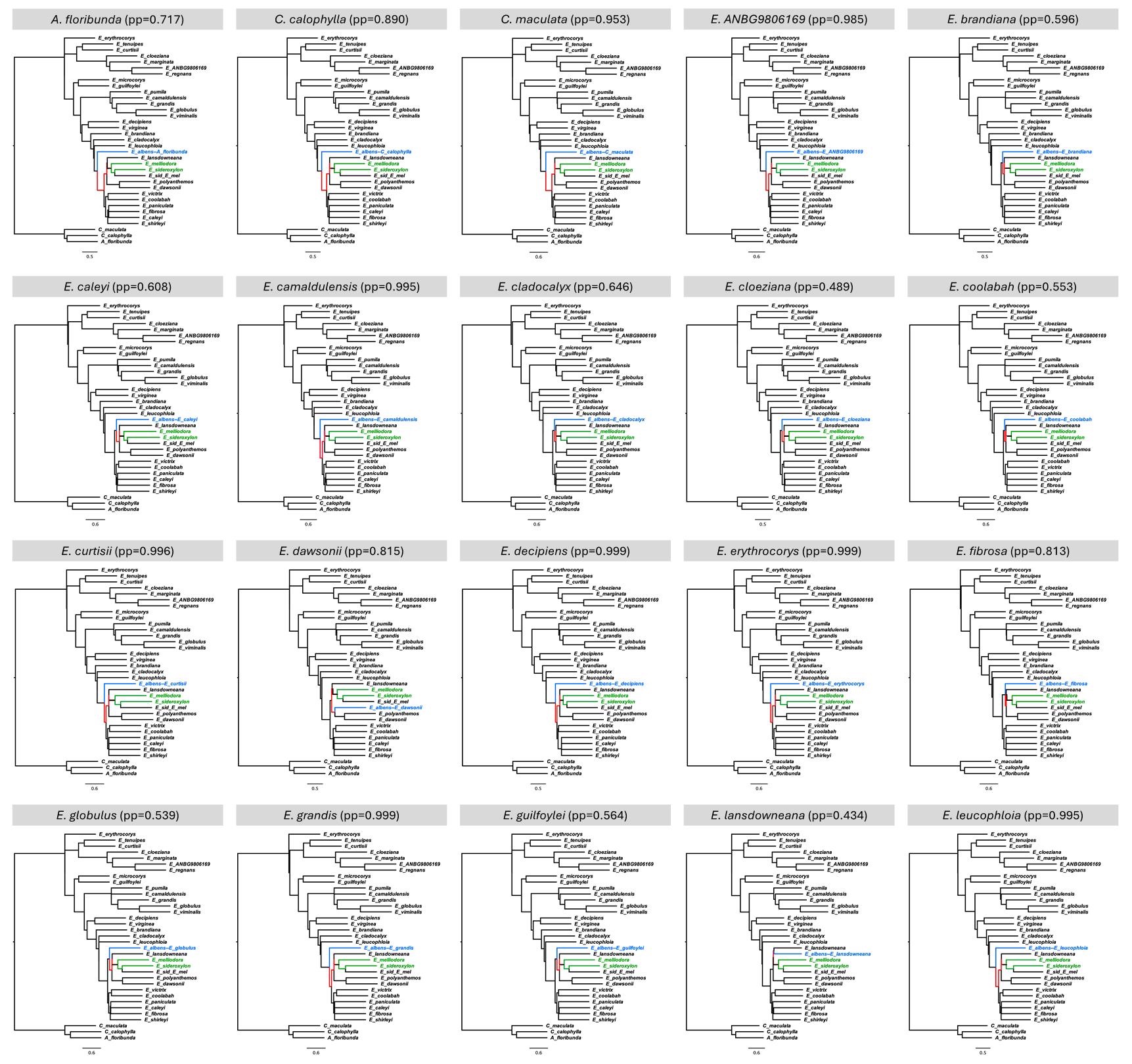

**Figure S15.** ASTRAL species trees for the short-read data of *Eucalyptus albens* across different reference genomes based on 112 BUSCO with non-significant results from both statistical tests (part 1). pp: ASTRAL posterior probability. Blue colour denotes *E. albens*; green colour denotes sister taxa of *E. albens* according to the published tree (Fig. S1); red colour denotes the positional shift of *E. albens* compared to the published tree (Fig. S1).

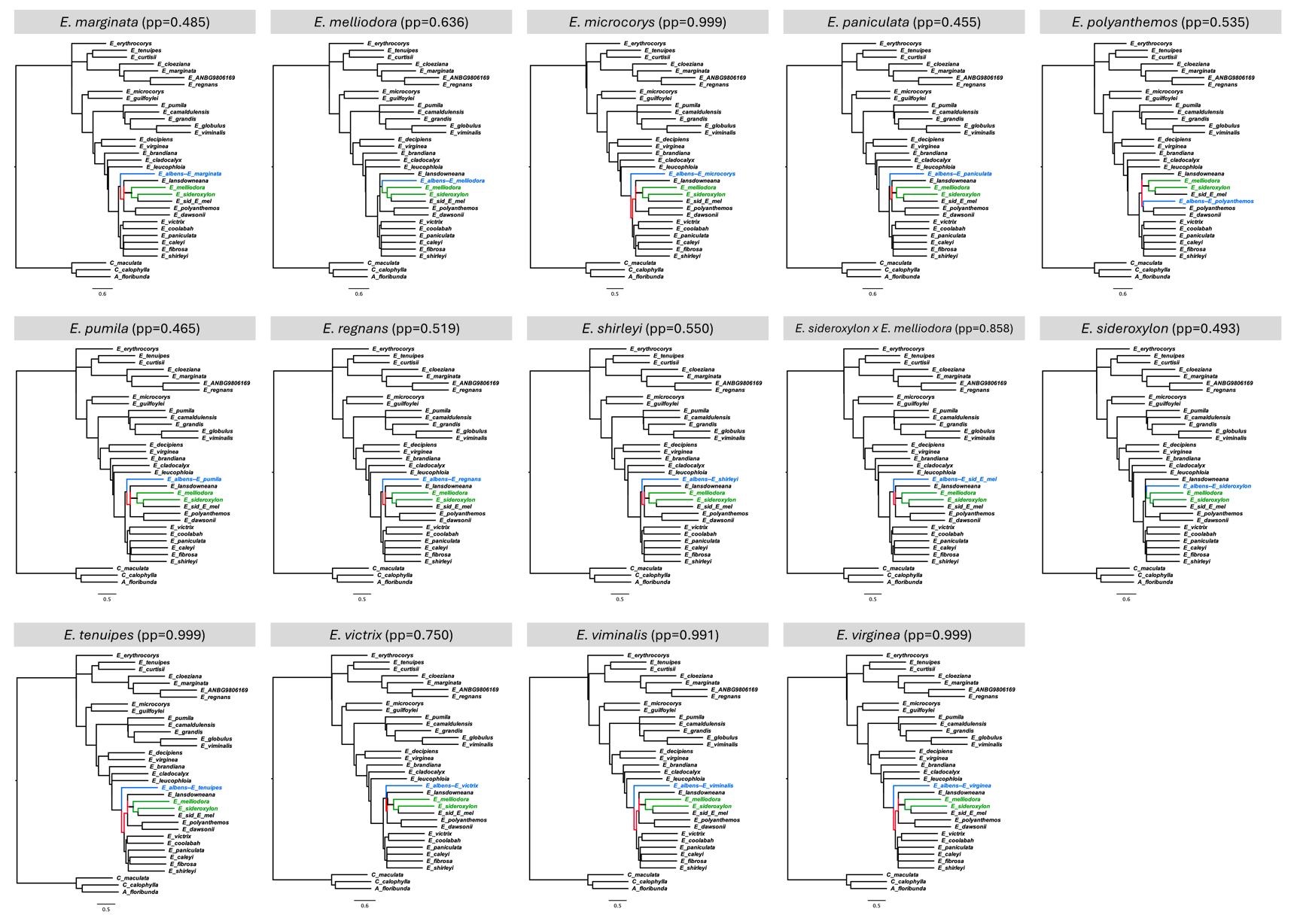

**Figure S16.** ASTRAL species trees for the short-read data of *Eucalyptus albens* across different reference genomes based on 112 BUSCO with non-significant results from both statistical tests (part 2). pp: ASTRAL posterior probability. Blue colour denotes *E. albens*; green colour denotes sister taxa of *E. albens* according to the published tree (Fig. S1); red colour denotes the positional shift of *E. albens* compared to the published tree (Fig. S1).

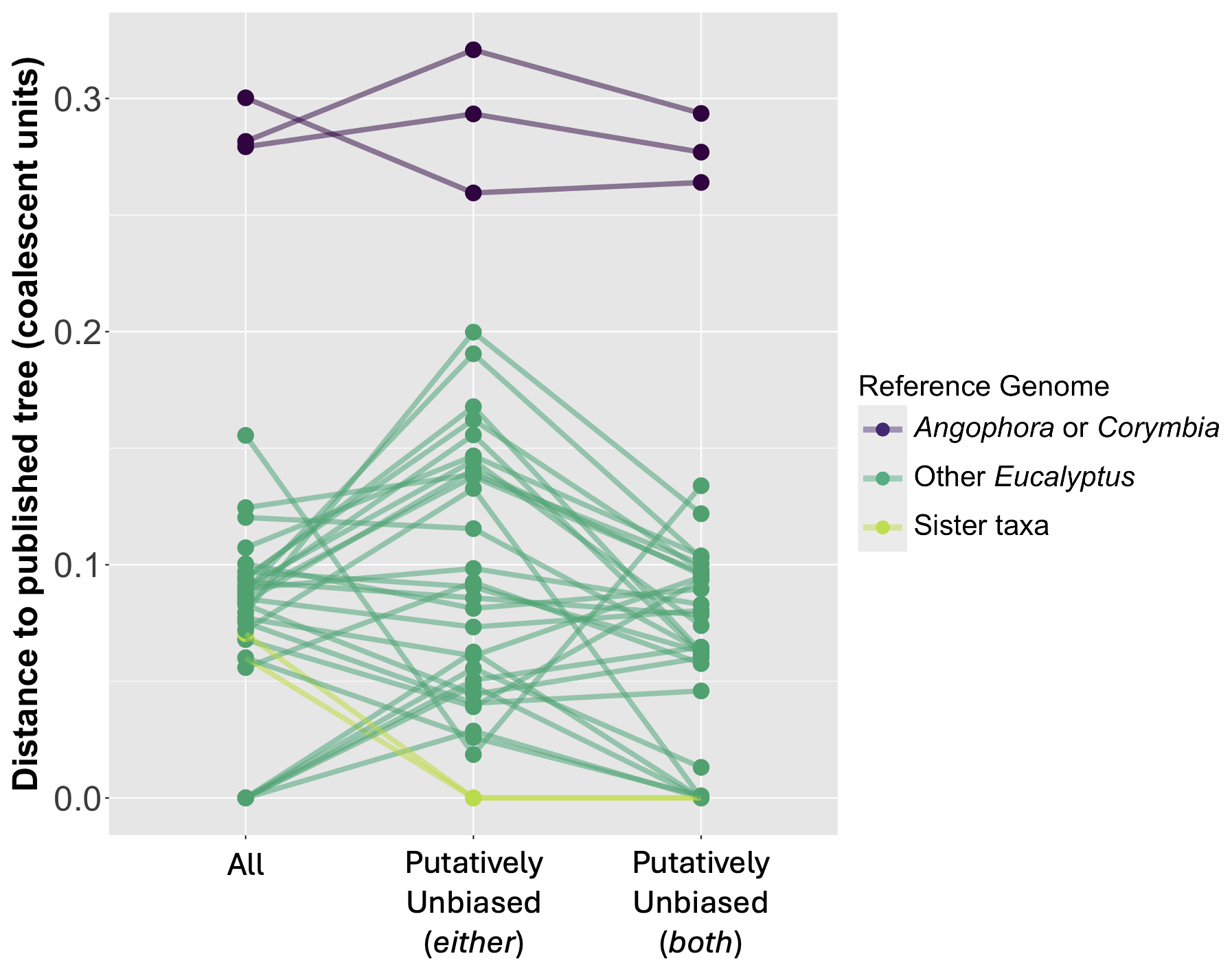

**Figure S17.** Phylogenetic distance between the position of *Eucalyptus albens* on the ASTRAL trees (inferred using *all* BUSCO loci, putatively unbiased loci based on *either* test, and putatively unbiased loci based on *both* tests) and the published tree (Ferguson et al. 2024a). The y-axis reflects the total branch lengths (in coalescent units) needed for the reconstruction to reach the ‘true’ branch on the published tree. Colouring denotes different reference genomes used for mapping (Fig. S1). Lines connect reconstructions with the same reference genome.
